# Ourotide: decoding the hierarchical peptide recognition for generative design

**DOI:** 10.64898/2026.09.10.750272

**Authors:** Yuhang Shen, Jie Zhang, Zhouyue Wu, Ziyan Xing, Qiuyu Yuan, Wen Zhang, Qigang Zhou, Feng Han, Nan Jiang, Xun Chen

## Abstract

The historical dichotomy between small-molecule pocket and extended protein interfaces misrepresents the physical reality of peptide recognition.^1,2^ Here we show that peptide binding is not a simple structural intermediate but a distinctly multimodal landscape comprising small-molecule-like pockets, protein-like interfaces, and a previously unrecognized third mode. This third regime is governed by a hierarchical subpocket architecture where a flattened surface achieves near-complete peptide engagement through spatially partitioned hydrophobic components. To maintain stability, an incompletely enclosed dominant anchor cooperates with highly hydrated auxiliary subpockets and an asymmetric receptor coupling mechanism that concentrates energy in an adjacent continuous water network. Because this unique binding mode suffers from extreme data scarcity, standard deep learning models fail to capture its physics.^3,4^ To resolve this, we mapped these specific geometric signatures to mine structurally faithful training distributions from global protein interactomes. Based on these data, we trained Ourotide, a deep learning framework coupling conditional geometric flow matching with interface-aware affinity learning. Evaluated across peptides up to 65 residues, Ourotide outperforms generalist models in backbone accuracy, interface recovery, and affinity prediction. This approach suggests that overcoming data scarcity in the physical sciences requires physics-guided data augmentation rather than naive statistical scaling.

## Introduction

Peptide therapeutics occupy a critical chemical space that diverges fundamentally from both traditional small molecules and proteins.^1,2,5,6^ Labeling them merely as structural intermediates offers no mechanistic blueprint for rational drug design. Categorizing these molecules by mass or residue count obscures a fundamental question about their binding modality. It remains unclear whether they bind within confined pockets like small molecules, sprawl across pre-organized interfaces like proteins or exploit unconventional recognition mechanisms. Without resolving this ambiguity, binding prediction remains empirical, leaving the principles that guide drug design undefined.^2,7^ This knowledge gap is particularly acute for flexible peptides. Their recognition relies neither on the rigid lock-and-key occupancy of small molecules nor the pre-folded docking of large proteins. Instead, it constitutes a complex thermodynamic negotiation among intermolecular contacts, conformational entropy, receptor plasticity, and solvation.^7–10^ Although unified generative models accommodate diverse molecular classes, a shared structural representation does not guarantee a shared physical understanding. The specific local and peripheral interactions required to simultaneously satisfy binding geometry and affinity for these flexible peptides remain elusive.^3,4,11,12^ Neural Networks has the potential to bridge this gap. However, indiscriminate data pooling masks the underlying physical principles governing the stabilization of distinct binding modes.^3,4,13,14^ To resolve this bottleneck for therapeutic peptides, we systematically compared 186 high-affinity-index peptide–receptor complexes with complexes formed by FDA-approved small molecules and with protein–protein interfaces. We reveal that the peptide binding space segregates into three distinct structural architectures, namely small-molecule-like pockets, protein-like interfaces, and a specialized hierarchical subpocket architecture.

This tripartite classification provides essential structural context. Existing drug or biologic prediction frameworks can increasingly address the first two modalities, but the hierarchical architecture remains a critical blind spot.^11,12,15,16^ This third modality suffers from extreme data scarcity and defies standard molecular paradigms.^17–20^ It features a flattened surface that maintains extensive peptide engagement through spatially separated hydrophobic subpockets. An incompletely enclosed dominant anchor is cooperatively stabilized by auxiliary subpockets and asymmetric receptor-shell coupling. This coupling is strongest immediately outside the anchor and fades towards the periphery alongside persistent cross-shell hydration.

Because existing models fail to capture these specific physical phenomena, we developed a targeted methodology explicitly for this data-scarce regime.^3,4^ Using these geometric and energetic signatures, we extracted hierarchical-peptide-like complexes from global protein-protein interactomes. This physics-guided augmentation preserves native site features that traditional solvent-accessible surface area heuristics fail to capture. Trained on this meticulously curated distribution, we introduce Ourotide. This generative framework couples conditional geometric flow matching with interface-aware affinity learning and complementary energy supervision. Under a best-of-five evaluation, Ourotide significantly improves both backbone and interface recovery for these uniquely challenging peptides while demonstrating superior affinity prediction. By isolating the most intractable mode of peptide recognition and anchoring generative models in its specific physical characteristics, this work provides a mechanistic foundation for next-generation peptide drug design. The study design and analytical workflow are summarized in Fig. 1.

**Fig. 1.**
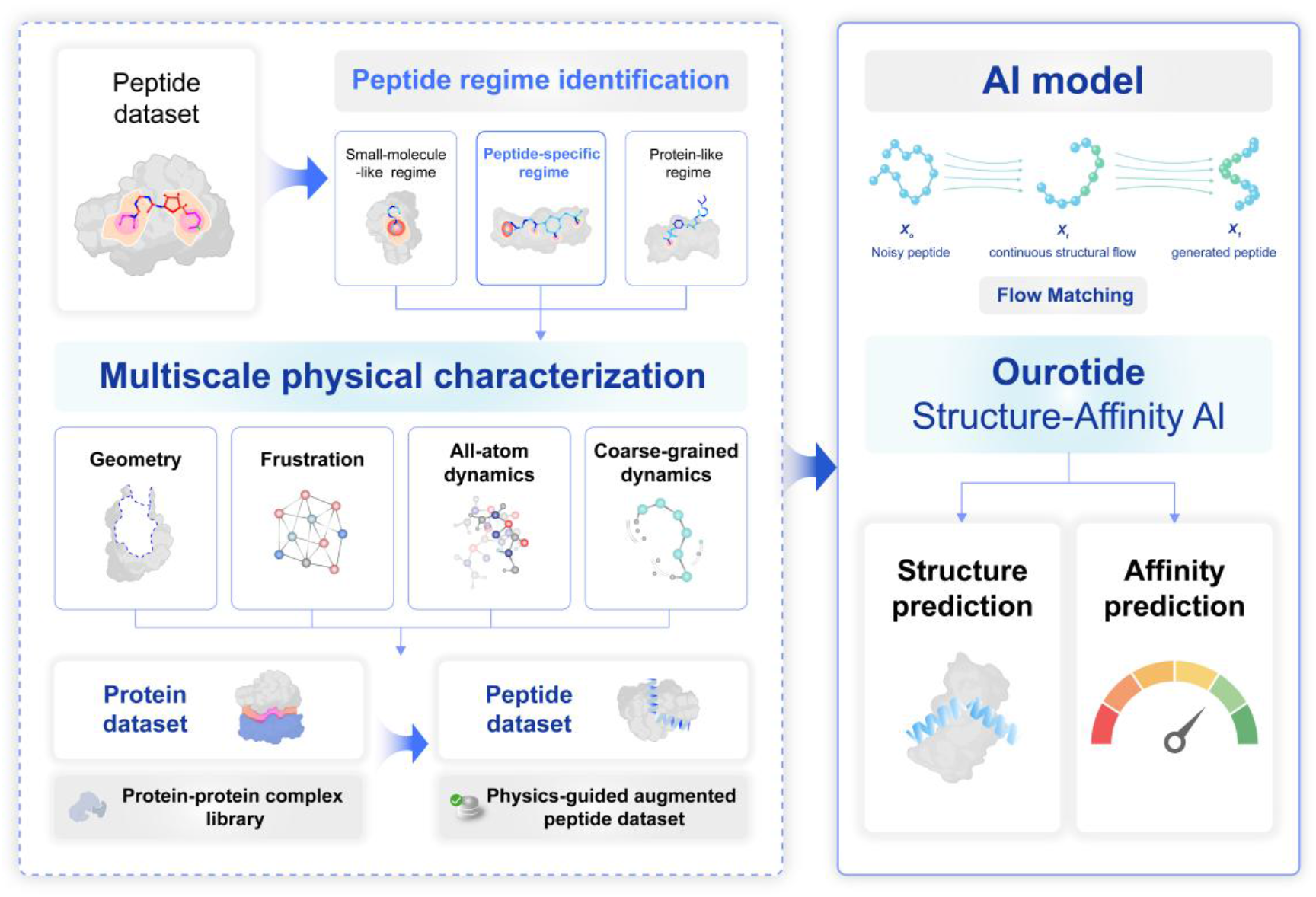
Overview of the physics-guided data strategy and Ourotide framework. Protein–peptide complexes were mapped into small-molecule-like, peptide-specific and protein-like recognition regimes. Geometry, configurational-frustration analysis, all-atom molecular dynamics and coarse-grained dynamics were used to define the physical features of peptide-specific recognition and to identify peptide-like receptor-binding segments in protein–protein complex data, yielding a physics-guided augmented peptide dataset. The augmented data were used to train the conditional Cα flow-matching structure generator and the interface-aware affinity predictor.

### Hierarchical Subpocket Architecture

Molecular recognition is traditionally anchored by two structural archetypes, which are compact pockets encapsulating small molecules and extended interfaces mediating protein interactions. Although peptide therapeutics bridge this size gap, their binding architectures defy a simple structural continuum. From a curated library of 186 structurally resolved high-affinity-index peptide–receptor complexes, we mapped binding sites onto a joint landscape defined by peptide contact fraction and the number of connected hydrophobic components (Fig. 2a). This landscape revealed three distinct density basins. Basin ii achieves extensive contact through a single hydrophobic core characteristic of small-molecule pockets. Basin iii degenerates into a protein-like interface mediating only restricted peptide segments. Conversely, basin i defines a peptide-specific regime characterized by a hierarchical subpocket architecture (Fig. 2a). While maintaining near-complete engagement, it strictly partitions hydrophobic interactions across multiple spatially isolated subpockets (Fig. 2b). This high-contact, distributed topological mode constitutes a structural anomaly and forms the central focus of our mechanistic analysis.

To determine whether basin i represents a mere geometric transition state or a novel binding regime with unique characteristics, we benchmarked these sites against canonical small-molecule pockets and protein interfaces. Geometrically, basin i occupies the intermediate space. It is flatter and thinner than small-molecule pockets yet thicker and less planar than protein interfaces (Fig. 2c,d). Topologically, it fractures this size-based continuum. Whereas unified single-component cores overwhelmingly dominate small-molecule pockets, and protein interfaces frequently disintegrate into higher-order fragments, basin i preferentially enriches a multi-component hydrophobic architecture (Fig. 2e). This frequency significantly surpasses both the small-molecule and protein reference groups . Thus, basin i occupies a demarcated topological regime that is more discrete than traditional pockets yet maintains a higher degree of spatial integration than extended interfaces.

Crucially, this structural fragmentation reflects a spatial reorganization rather than a decrease of hydrophobic residues. Total hydrophobic fractions overlap broadly across all three modalities. The basin i distribution exhibits two density maxima that precisely capture the physicochemical environments of both canonical extremes. Basin i is not a simple average of pockets and interfaces. Instead, it operates as a chimeric recognition mode where multiple interface-like, spatially discontinuous hydrophobic patches synergistically assemble to achieve a pocket-like, high-contact engagement (Fig. 2f). Therefore, peptide recognition reveals a distinctly multimodal landscape where basin ii replicates the pocket feature, basin iii mirrors the localized interface, and basin i fractures and recombines both structural characteristic.

**Fig. 2.**
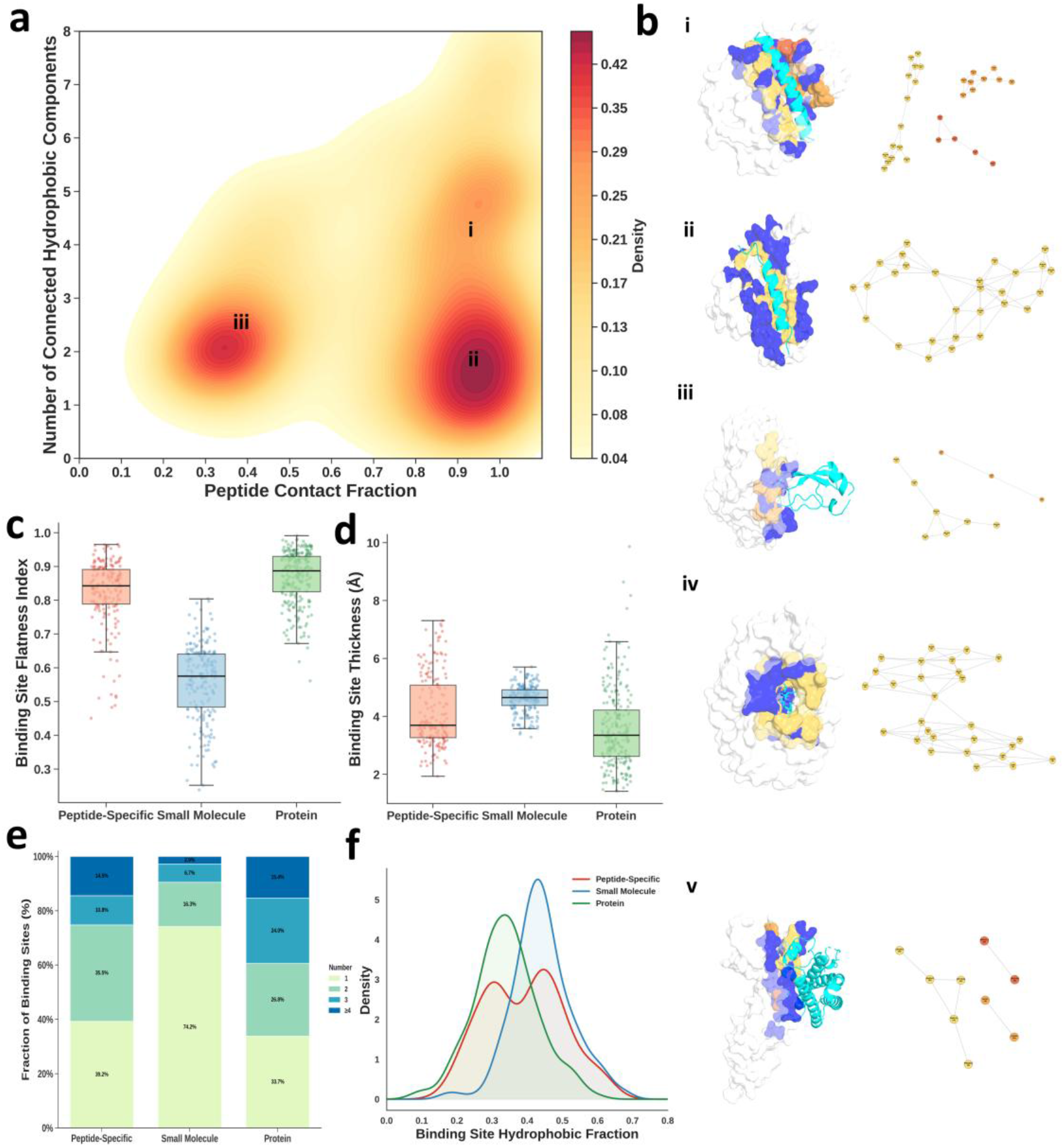
Geometric characteristics and hydrophobic partitioning of peptide-binding sites. a, Joint density map of peptide contact fraction and the number of connected hydrophobic components across 186 high-affinity-index protein–peptide complexes. Three high-density regions are annotated as i, ii, and iii. b, Representative complex structures and their corresponding hydrophobic connectivity graphs are shown on the right. In the structural models (PDB IDs 1DPJ, 1G5J, and 2PTC), hydrophobic binding-site residues belonging to the largest (C1), second-largest (C2) and third-largest (C3) connected components are coloured yellow, orange and red, respectively; hydrophilic binding-site residues are coloured blue, and peptide ligands are shown in cyan. The same colour scheme is used in the connectivity graphs, in which nodes represent hydrophobic receptor residues, nodes of the same colour belong to the same connected component, and edges denote pairwise connections. c,d Box-and-point distributions of binding site flatness index (c) and thickness (d) for peptide-binding sites (red), small-molecule pockets(blue), and protein-protein interfaces (green). e, Proportions of binding sites containing one, two, three, or at least four connected hydrophobic components across the three structural modalities. f, Density distributions of the total binding site hydrophobic fraction. Colors match those in panel b. Representative small-molecule (PDB ID 1MUI, structure notated iv) and protein interface (PDB ID 1A22, structure notated v) complex structures are shown alongside.

### Hierarchical Subpocket Recognition

To determine whether the hierarchical subpocket architecture is merely a collection of miniature drug-like pockets, we compared the largest peptide component to small-molecule pockets and protein interfaces. Contact-averaged frustration placed this dominant anchor directly between highly optimized small-molecule pockets and diffuse protein interfaces (Fig. 3a). Representative structures explain this intermediate position. Stabilizing contacts line opposing walls to envelop a small molecule, whereas they concentrate on just one face of the bound peptide and remain diffuse across protein interfaces (Fig. 3b). The dominant anchor therefore retains a focal stabilized core but lacks the surrounding reinforcement of a closed cavity.

This structural incompleteness directly limits dynamic confinement. Across molecular dynamics trajectories, geometric enclosure decreased sharply from small-molecule pockets to peptide subpockets, and finally to open protein interfaces (Fig. 3c). Protein interfaces were further distinguished by large interfacial gaps and pervasive hydration, whereas the peptide and small-molecule sites remained structurally proximate (Fig. 3d,e). Thus, the dominant anchor preserves a stable local foothold but cannot provide the full geometric enclosure characteristic of a canonical drug pocket.

Because this dominant anchor is structurally incomplete, auxiliary subpockets must reinforce it. This reinforcement relies on a defined energetic hierarchy rather than a set of equivalent pockets. Local stabilization decreases sequentially from the dominant anchor to the auxiliary subpockets (Fig. 3f). The physical necessity of this hierarchy is evident in the free-energy landscape. The absolute free energy minimum requires the dominant anchor to engage alongside the auxiliary subpockets. Partially engaged states remain at higher free energies even when the dominant anchor is perfectly retained (Fig. 3g, 3h).

These auxiliary subpockets are not merely smaller copies of the rigid anchor. In contrast, they are significantly less enclosed and more persistently hydrated (Fig. 3i, 3j). Rather than forming secondary rigid cavities, they function as a plastic, solvent-accessible network that extends stabilization across the peptide surface (Fig. 3k). Small molecules concentrate stabilization within one closed pocket while proteins distribute it broadly. Hierarchical subpocket recognition adopts a distinct, asymmetric solution. It utilizes subpocket multiplicity not to eliminate energetic hierarchy, but to stabilize a broad binding surface that no single local pocket can fully enclose.

**Fig. 3.**
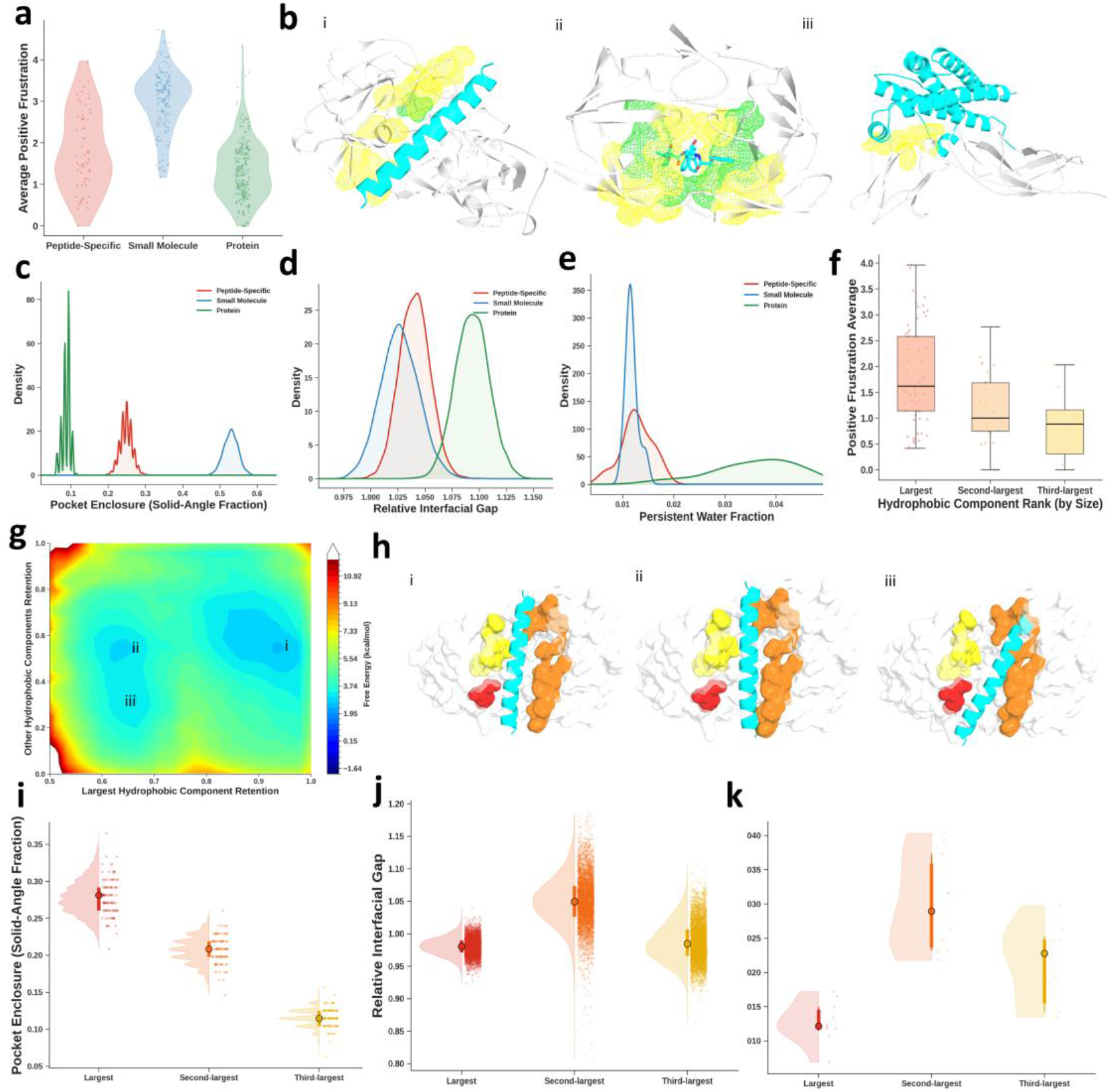
Hierarchical subpockets distribute stabilization across peptide-specific sites. a, Average positive-frustration scores for the largest connected hydrophobic component. b, Representative structures contrast the single-face, concentrated stabilization of peptides with the enclosing small-molecule cavities and diffuse protein interfaces (PDB IDs 1DPJ, 1MUI, and 1A22). Peptide, small-molecule, and protein sites are colored red, blue, and green, respectively. The largest three peptide subpockets (C1, C2, and C3) are colored red, orange, and yellow. Receptors are white, ligands are cyan, and highly or minimally frustrated contacts are yellow or green. c,d,e, Molecular dynamics-derived distributions of pocket enclosure (solid-angle fraction, c), relative interfacial gap (d), and persistent-water fraction (e). f, Average relative-frustration scores for C1, C2, and C3 (three hydrophobic component of hierarchical subpocket peptide). g, Two-dimensional free-energy surface projected onto the retention of C1 versus the remaining components. h. Corresponding structural snapshots are shown on the right. The largest three peptide subpockets (C1, C2, and C3) are colored red, orange, and yellow. Receptors are white, ligands are cyan. i,j,k Component-resolved structural dynamics including enclosure (i), interfacial gap (j), and persistent hydration (k).

### Asymmetric Shell Cooperativity

The hierarchical subpocket architecture, characterized by hydrophobic components interspersed with hydrophilic residues, necessitates a stabilization mechanism that extends beyond its discontinuous local core. Mapping energetic frustration radially outward from the binding pocket (S0) reveals that peptide stabilization relies on a spatially extended, monotonic gradient (Fig. 4a–c). Unlike Small-molecule sites abruptly lose stabilization outside their optimized pocket (Fig. 4b), and protein interfaces display non-hierarchical energetic topographies (Fig. 4c), peptide sites gradually redistribute their stabilizing burden across successive peripheral shells (S1 to S3, Fig. 4a) . This topological discontinuity does not induce structural chaos. Rather, it translates into a highly coordinated dynamic response (Fig. 4d). Distance-fluctuation maps demonstrate that while small-molecule pocket decouple from their surroundings, peptide complexes propagate structured, high-fluctuation bands across the S0 to S2 boundaries (Fig. 4e). Consequently, the local geometric weakness of a fragmented pocket is offset by recruiting the surrounding receptor into a broadly adaptable and directionally organized dynamic network.

Projection of the free-energy landscape exposes a thermodynamic asymmetry governing this extended network (Fig. 4f). Maximal stability (state i) dictates the concurrent maintenance of contacts across multiple shell boundaries, yet these interfaces contribute unequally to the global minimum. Disruption of the outer S1 to S2 network (state ii) exacts only a minor thermodynamic penalty, whereas uncoupling the inner S0 to S1 boundary drives severe destabilization (Fig. 4g). This energetic topology identifies the immediately adjacent shell (S1) as a critical stabilizing collar. It energetically compensates for the hydrophilic interruptions that physically divide the primary core. This collar couples strongly to the anchor while interacting with distal shells through progressively attenuated forces. The peptide binding site therefore establishes a soft cooperativity. It functions neither as a uniformly rigid lock nor a completely fluid interface, but as a structurally tiered system that concentrates stabilization energy immediately outside the discontinuous primary pocket.

This asymmetric shell coupling is physically mediated by a distinctive and continuous hydration architecture. Persistent water concentrates heavily within the S1 collar, effectively wetting the hydrophilic residues that partition the hydrophobic subpockets (Fig. 4h). However, the defining signature of peptide recognition emerges further outward. As the core water cloud relaxes from a highly anisotropic geometry toward an isotropic distribution in the periphery, the cross-shell connectivity of shared waters remains remarkably unbroken from S0 to S3 (Fig. 4i,4j).

This persistent solvent matrix evades the rapid radial decay typical of both small-molecule and protein interfaces. Instead, it acts as a continuous fluid scaffold that bridges the physically separated subpockets and sustains weak inter-shell communication. Ultimately, peptide binding operates through a nested radial hierarchy. This includes a focal but physically interrupted anchor (S0), a strongly coupled hydration collar (S1), and a compliant peripheral support network (S2 and S3). This tiered architecture seamlessly resolves the intrinsic instability of fragmented pockets, achieving global structural integrity while preserving the local adaptability essential for peptide recognition.

**Fig. 4.**
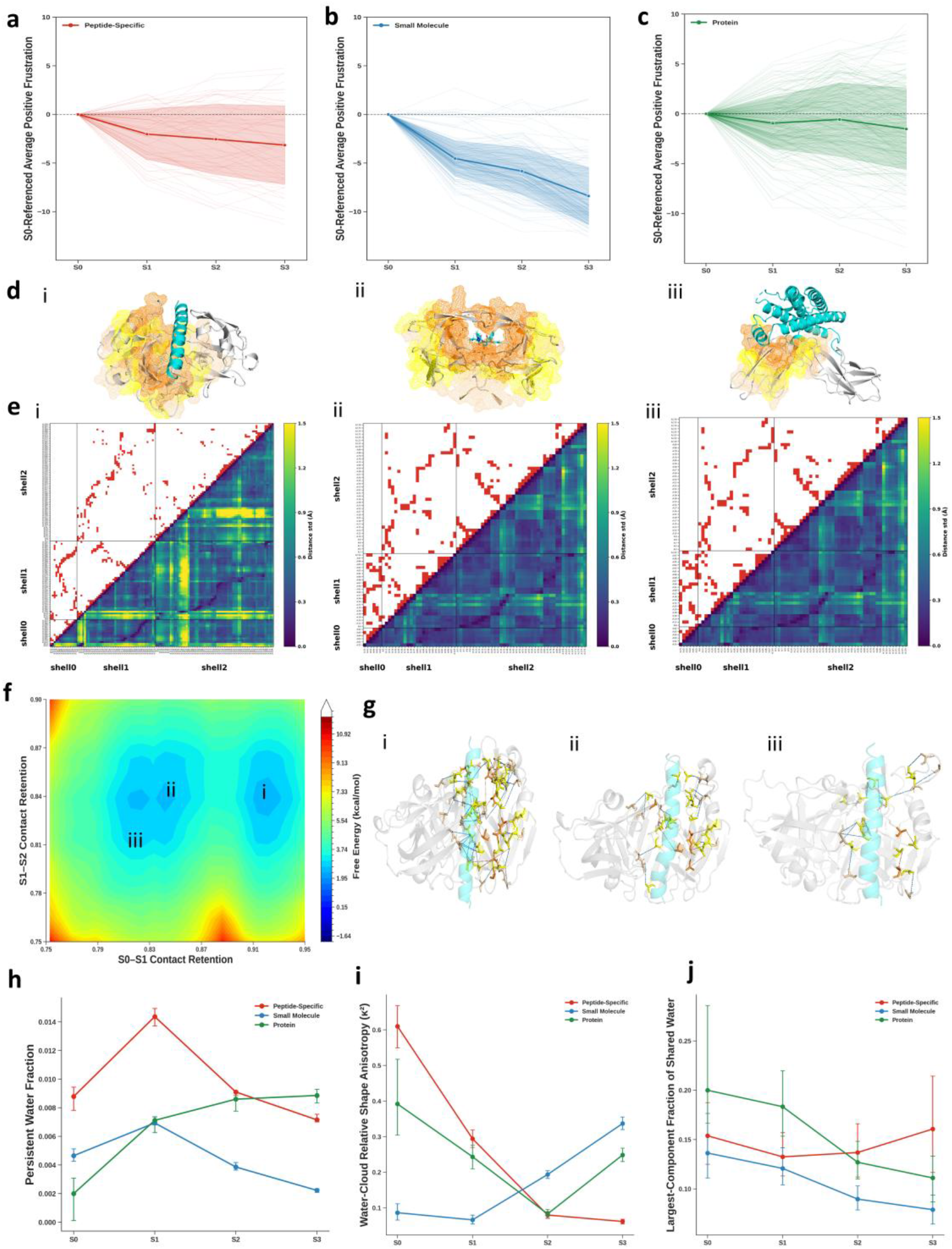
Shell-resolved energetic, dynamic, and hydration architecture of peptide binding. a,b,c Shell-resolved profiles of average positive frustration referenced to the pocket (S0) and successive peripheral shells (S1 to S3) for hierarchical subpocket peptide (a), small molecule pocket (b) and interface (c). Thin curves denote individual sites, and bold curves denote ensemble averages. d, Bottom structures illustrate the corresponding cross-shell spatial organization (PDB IDs 1DPJ, 1MUI, and 1A22). Receptors are shown as white cartoons, and peptide or protein ligands are shown as cyan cartoons; the small-molecule ligand is shown as cyan sticks. Receptor shells S1, S2 and S3 are displayed as orange, yellow and wheat-coloured mesh surfaces, respectively. e, Combined residue-contact and distance-fluctuation maps for representative complexes. Upper triangles display residue contacts as red pixels. Lower triangles map the standard deviation of inter-residue distances in Angstroms via a color scale. Gray lines demarcate the S0 to S2 shell boundaries. f, Free-energy surface of a representative basin i peptide complex projected onto S0 to S1 and S1 to S2 contact-retention coordinates. g, Extracted structures (states i, ii, and iii) illustrate distinct cross-shell contact configurations along the energy landscape. h,i,j Shell-resolved hydration properties spanning S0 to S3. Metrics comprise the persistent-water fraction (h), water-cloud relative shape anisotropy (i), and the largest-connected-component fraction of adjacent-shell shared waters (j).

### Hierarchy-aware peptide prediction

Given the scarcity of structural data, robust data augmentation is essential for generative peptide drug design. However, conventional augmentation based on solvent-accessible surface area heuristics fails to recapitulate the hierarchical complexity of native peptide-binding sites. These generated sites are predominantly single-component and less flat (Fig. 5b,c). They lack the distinct frustration contrasts and multi-component organization unique to genuine peptide interfaces (Fig. 5d,e). In contrast, we targeted complexes based on hydrophobic signatures, flatness, and shell-level energetics characteristic of true binding basins (Fig. 2a). This aligned our training distribution with the physical reality of hierarchical subpockets.

Using the physics-guided dataset, we developed Ourotide as a framework comprising a conditional Cα flow-matching structure generator and a trained interface-aware affinity predictor (Fig. 5f). Ourotide covers peptides spanning 8–65 residues, extending beyond the supported length range of several peptide-specific comparators. We therefore evaluated five methods on a strict common set of 138 complexes spanning the full length range and seven methods on a strict common set of 47 complexes within the shared shorter-peptide range. When evaluating the best of five predictions for each method, Ourotide recovered the lowest-Cα-RMSD structures for a larger fraction of targets than all applicable comparators, including AlphaFold 3 (Fig. 5g, 5h).^11,21–26^ Thus, this broader length coverage was accompanied by improved best-sampled reconstructions on targets accessible to all baseline methods. Furthermore, these same RMSD-selected structures achieved a higher overall Q_interface (Fig. 5i, 5j), demonstrating that this improvement extended beyond mere backbone placement to the accurate recovery of native receptor-peptide interactions.

While AlphaFold 3 also accommodated these longer chains, it did not reconstruct the hierarchical interfaces as faithfully.^11^ Specifically, Ourotide reproduced flattened surface binding in 4DRA (Fig. 5k, i) and the simultaneous occupancy of separated hydrophobic subpockets in 4AYE (Fig. 5k, ii). It also preserved distributed contacts in 3IA3 (Fig. 5k, iii) and an elongated, multi-region binding footprint in 2UXN (Fig. 5k, iv). Across these examples, Ourotide’s residue-pair maps more closely reproduced the native interaction patterns than those of AlphaFold 3.

In addition to structure generation, the trained interface-aware module provided quantitative affinity predictions for the same class of receptor–peptide complexes. Ourotide achieved the highest affinity-prediction success rates among seven evaluated methods, with over 81.6% of predictions falling within |ΔpKd| ≤ 2.0 (Fig. 5l).^21,27–31^ In ablation experiments, replacing the physics-guided augmented set with the geometrically augmented set reduced both structure-recovery and affinity-prediction performance, supporting a contribution from the physically filtered training distribution. Ultimately, the binding hierarchy is more than a descriptive structural classification; by embedding it into the augmentation strategy, Ourotide forces the generative process to respect the physical constraints of the pocket. This alignment yields measurable gains in both structural fidelity and affinity, establishing a computational framework for generating peptide therapeutics against highly structured, hierarchical targets.

**Fig. 5.**
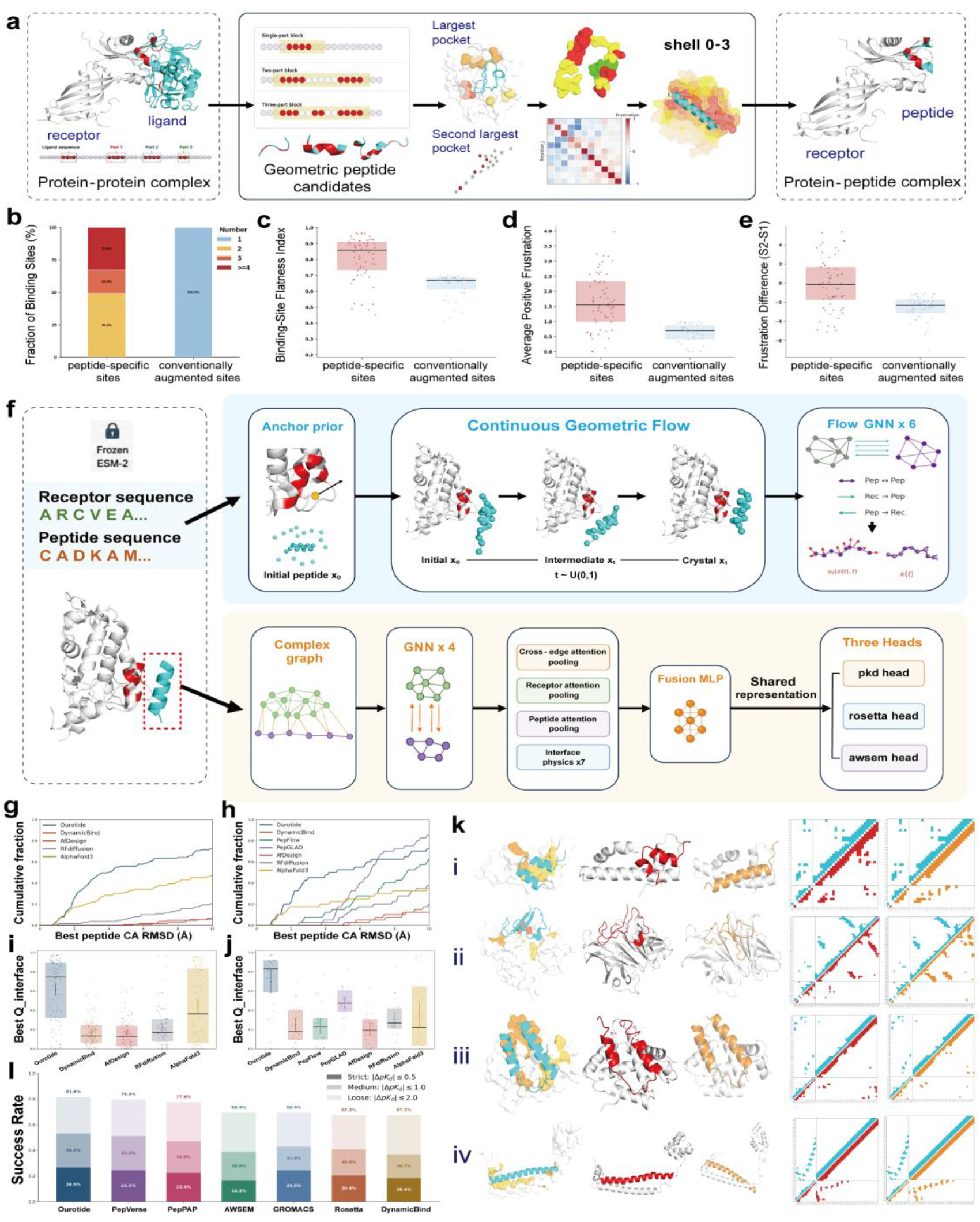
Physics-guided flow matching for peptide–receptor structure and affinity modeling. a, Generation of physics-augmented peptide-receptor complexes from protein-protein interfaces. Candidate geometries are filtered by hydrophobic organization, site flatness, subpocket stabilization, and shell energetics. Receptors (white or light gray) and interacting chains (cyan) are shown with colored patches denoting hydrophobic subpockets. b,c,d,e Structural and energetic comparison of peptide-specific versus solvent-accessible surface area-derived binding sites. Features include hydrophobic-component count (b), site flatness (c), mean positive frustration in the largest hydrophobic cluster (d), and the frustration difference between shell 2 and shell 1 (e). Stacked bars denote component counts as blue (1), yellow (2), orange (3), and red (4 or more). f, Ourotide deep learning architecture. A hotspot-guided conditional C-alpha geometric flow-matching module generates peptide conformations. g,h Cumulative distribution of best-of-five peptide C-alpha root-mean-square deviation for the full-length (up to 65 residues, g) and shorter-peptide benchmarks (h). i,j Interface recovery metric evaluated on the lowest-RMSD models. The left panel shows the five-method full-length benchmark (j), and the right panel shows the seven-method common benchmark(i). k, Representative binding architectures (PDB IDs 4DRA, 4AYE, 3IA3, and 2UXN) comparing experimental structures (left, cyan), Ourotide predictions (middle, red), and AlphaFold 3 predictions (right, orange). Receptors are white or light gray with highlighted hydrophobic subpockets in yellow or orange. Lower panels display corresponding residue-pair contact maps. l, Affinity prediction performance across seven methods evaluated at strict, intermediate, and permissive error thresholds. Lighter shades represent cumulative model inclusion at relaxed thresholds.

## Discussion

Molecular recognition likely operates along a continuous physical spectrum governed by a universal set of interactions.^32–34^ Our findings suggest that peptide binding spans this entire continuum. Besides some peptides bind via single cavities reminiscent of small molecules and others utilize extended protein-like interfaces, the most distinguishing feature of the third type appears to be a hierarchical subpocket architecture. Historically, drug discovery has segregated small molecules, peptides, and proteins into isolated therapeutic silos.^1,5,6^ However, this categorization may not reflect rigid structural boundaries. It is more likely an artifact of historical scale effects driven by past limitations in chemical synthesis and experimental accessibility. Consequently, we propose that molecular modalities might be better understood as connected regimes across a shared landscape rather than discrete classes defined by historical constraints. This historical segregation highlights a potential limitation of current AI approaches in the physical sciences. In fields with abundant human-generated data such as natural language processing, empirical scaling laws have driven considerable success.^35^ However, these paradigms often face challenges at scientific frontiers where high-quality observations remain sparse. Increasing model capacity on datasets skewed by historical synthesis biases risks overfitting to these artifacts.^13,14,36^ A highly expressive model may struggle to recover structural principles that are underrepresented in the training set, rendering naive statistical data augmentation less effective in this context. We sought to address this data scarcity through targeted, mechanism-guided extraction rather than arbitrary augmentation. By hypothesizing that the geometric characteristics defining peptide recognition are preserved in data-rich protein-protein interfaces, we augmented the sparse target distribution using consistent structural metrics. This approach suggests that a curated set of mechanistically coherent examples could be more informative than larger, structurally confounded datasets. This physics-guided strategy for reconstructing data-limited learning distributions is summarized in Fig. 6.

**Fig. 6.**
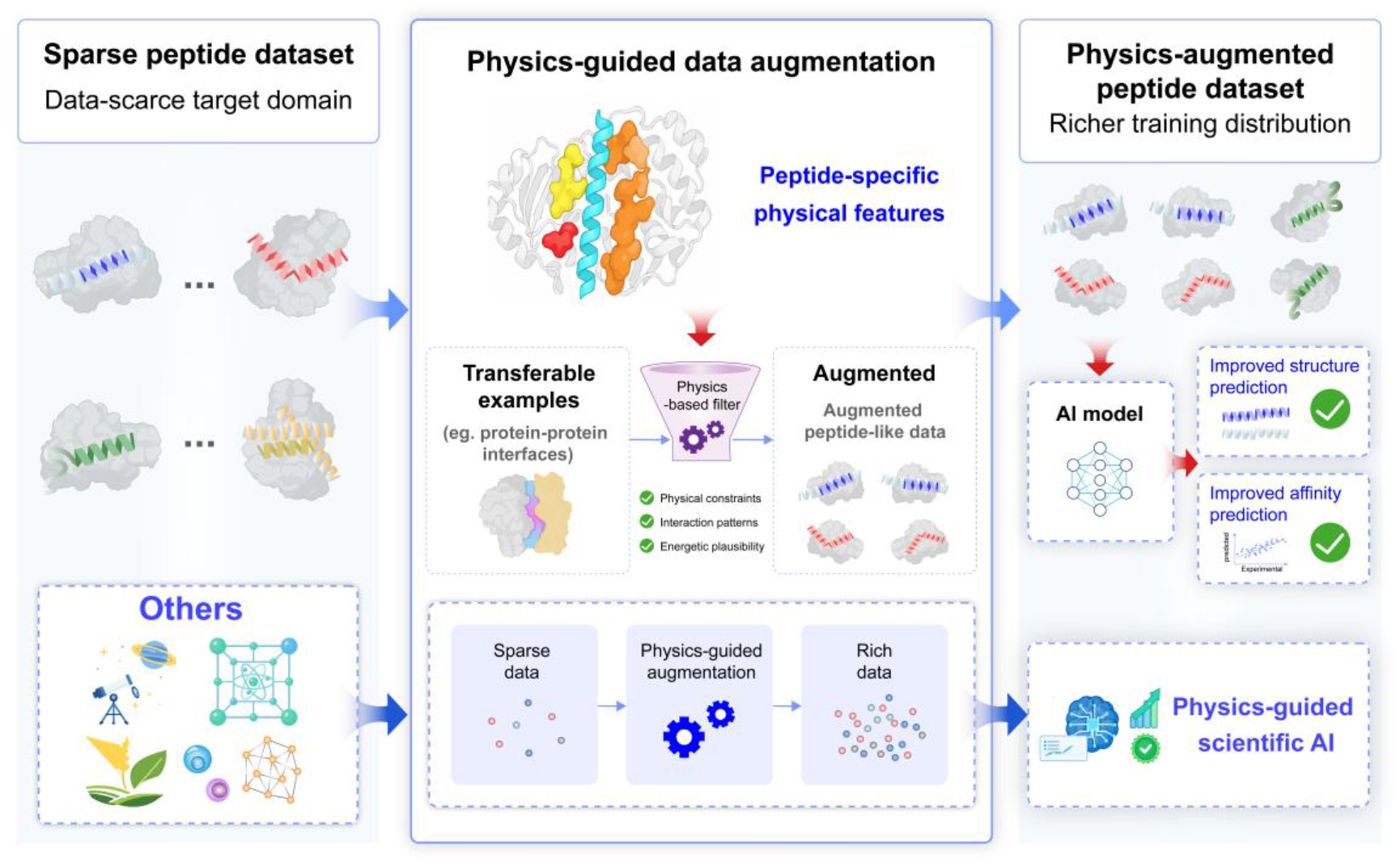
Physics-guided reconstruction of data-limited learning distributions. In data-scarce target domains such as peptide recognition, physically transferable examples can be identified in neighbouring data-rich domains and filtered using target-relevant structural, interaction and energetic criteria. For peptide recognition, protein–protein interfaces provide candidate receptor-binding segments that can be filtered according to peptide-specific physical features to construct an augmented peptide dataset. The resulting distribution supports improved structure and affinity prediction. More generally, the framework illustrates how physical knowledge can guide data selection and transfer in scientific domains where direct observations remain limited.

Together, these observations point toward a unified framework for AI modeling in data-limited scientific settings.^37^ While massive datasets can reveal underlying structures in data-rich domains, domain expertise must actively shape the learning problem where observations are scarce. This involves defining the problem through consistent features, transferring relevant data from neighboring domains, and reserving costly experiments for uncharted boundaries. While model capacity remains important, it cannot alone determine whether a training distribution reflects scientific reality. Ultimately, progress in AI for the physical sciences will likely depend on engineering learning distributions that faithfully capture the underlying physical phenomena.

## Methods

### Reference complex libraries

Protein complexes containing small-molecule, peptide or protein ligands were collected from PDBbind 2020.^38^ Structures with a PDBbind affinity index greater than 7, derived from reported Kd, Ki or IC50 measurements, were retained. Small-molecule complexes were further cross-referenced against a curated list of FDA-approved drugs, yielding approximately 387 protein–small-molecule complexes.

For complexes containing two standard amino-acid chains, the shorter and longer chains were designated the ligand and receptor, respectively. Ligands of 8–65 residues defined the protein–peptide library (n = 186), whereas ligands longer than 65 residues defined the protein–protein reference library (n = 246). FDA approval status was not used to select either library.

A receptor residue was assigned to the binding site when any of its atoms was within 6.5 Å of the ligand. Ala, Val, Leu, Ile, Met, Phe, Trp and Pro were treated as hydrophobic. Hydrophobic binding-site residues were represented as graph nodes and connected when their Cα atoms were within 9.5 Å. Peptide engagement was defined as the fraction of peptide residues containing at least one atom within 6.5 Å of the receptor. Complexes with peptide engagement below 0.5 were classified as protein-like. Among complexes with engagement between 0.7 and 1.0, those containing one hydrophobic component were classified as small-molecule-like, whereas those containing at least two components were classified as peptide-specific. Complexes outside these core regions were excluded from regime-specific analyses.

### Binding-site analyses

The binding-site hydrophobic fraction was calculated as the number of hydrophobic residues divided by the total number of binding-site residues. Principal-component analysis was performed on centred binding-site Cα coordinates. With covariance eigenvalues ordered as λ_1_ ≥ λ_2_ ≥ λ_3_, site flatness and thickness were defined as

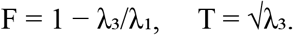

Pocket enclosure was calculated from molecular-dynamics trajectories as the fraction of 96 directions sampled on a unit sphere centred at the ligand centre of mass that were occluded by binding-site atoms.

The relative interfacial gap was calculated as the mean receptor–ligand surface separation in each frame normalized by its value in the reference structure.

Water-residence events were identified using entry and exit cutoffs of 0.35 and 0.40 nm, respectively. Persistent hydration was defined as the fraction of events lasting at least 500 ps. Trajectories were sampled every 50 ps and analysed in 25-ns windows.

The largest connected hydrophobic component was designated S0. Shell S1 comprised receptor residues containing a heavy atom within 4.5 Å of S0. Shells S2 and S3 were generated recursively using the same cutoff while excluding residues assigned to inner shells. Shell-only waters were within 0.35 nm of a shell heavy atom and more than 0.35 nm from every ligand heavy atom. Water-cloud relative shape anisotropy was calculated as

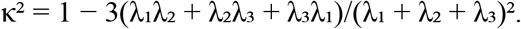

Waters within 0.35 nm of two adjacent shells were classified as shared. Shared-water networks used water oxygen atoms as nodes and a 0.35-nm distance cutoff for edges. Connectivity was quantified as the fraction of shared waters in the largest connected component. S0 and S3 were represented by the S0–S1 and S2–S3 shell pairs, respectively, whereas S1 and S2 were represented by the mean of their two adjacent shell pairs.

### Configurational-frustration analysis

Local configurational frustration was calculated for receptor structures using Frustratometer v0.3.2 and the AWSEM energy model.^28,39^ For each native intrareceptor contact (i,j), the configurational-frustration index was

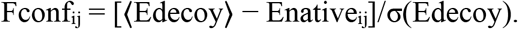

Here, Enativeᵢⱼ is the native AWSEM contact energy, and ⟨Edecoy⟩ and σ(Edecoy) are the mean and standard deviation of 4,000 configurational-decoy energies. Larger positive values indicate contacts that are energetically more favourable than their decoys.

For each residue, the positive configurational-frustration score was

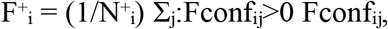

where N⁺ᵢ is the number of positive-index contacts involving residue i. Component- and shell-level scores were obtained by averaging F⁺ᵢ over residues in the corresponding region. These quantities describe intrareceptor energetic organization and were not interpreted as direct experimental binding free energies.

### Molecular-dynamics simulations

All-atom simulations were performed using GROMACS 2024.5^29^ for representative small-molecule, peptide and protein complexes 1MUI, 1DPJ and 1A22, respectively. Systems were prepared with CHARMM-GUI v3.7^40^ in periodic cubic boxes containing CHARMM-modified TIP3P water^41^ and 0.15 M NaCl. Proteins and peptides used CHARMM36m^42^, and the 1MUI ligand was parameterized with the CHARMM General Force Field.^43^

Following position-restrained steepest-descent minimization, each system underwent 125 ps of NVT equilibration with a 1-fs timestep. Protein or peptide backbone and side-chain heavy atoms were restrained using force constants of 400 and 40 kJ mol^−1^ nm^−2^, respectively; small-molecule heavy atoms used 400 kJ mol^−1^ nm^−2^.

Unrestrained 500-ns NPT production simulations were performed at 300 K and 1 bar with a 2-fs timestep, using the velocity-rescaling thermostat^44^ and C-rescale barostat.^45^ Electrostatics were treated with particle-mesh Ewald^46^ and a 1.2-nm real-space cutoff. Van der Waals interactions were force-switched between 1.0 and 1.2 nm, and hydrogen-containing bonds were constrained with LINCS.^47^ Five independent trajectories were generated per system using different initial velocities. Analyses used the final 200 ns of each trajectory.

### AWSEM free-energy calculations

Umbrella sampling was performed for the representative peptide–receptor complex 1DPJ using OpenAWSEM with the standard AWSEM potential.^28,48^ The collective variable was the native receptor–peptide interfacial-contact fraction, Q_int_. The native contact set I contained receptor-chain A and peptide-chain B residue pairs whose Cα atoms were within 0.95 nm in the reference structure:

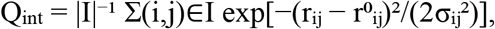

where rᵢⱼ and r⁰ᵢⱼ are the instantaneous and reference distances, respectively, and

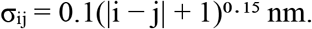

Here, i and j denote the zero-based sequential residue indices within receptor chain A and peptide chain B, respectively, following the implementation used to construct the collective variable.

A harmonic bias was applied along Q_int_:

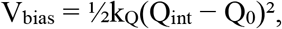

with k_Q_ = 100 kcal mol^−1^. Twenty-one windows spanning Q_0_ = 0–1 at intervals of 0.05 were each simulated for 2.0 × 10^6^ AWSEM steps at 300 K.

Two protocols were performed. In the first, neither chain received additional positional restraints. In the second, receptor chain A was restrained to its reference coordinates using a force constant of 10,000 kcal mol^−1^ nm^−2^, while peptide chain B remained mobile. Relative free-energy profiles were reconstructed using the weighted histogram analysis method^49^, and two-dimensional landscapes were obtained by projecting the WHAM-reweighted ensembles onto the indicated structural coordinates.

### Protein-derived peptide library

Both chain orientations of each protein–protein complex were evaluated by alternately assigning one chain as the receptor and the other as the source of candidate peptide segments. Receptors were required to contain at least 66 standard amino-acid residues. A source-chain residue was classified as interfacial when it formed at least three heavy-atom pairs within 4.5 Å of the receptor. Contact residues separated by no more than three sequence positions were grouped into continuous segments. Up to three segments were combined when intervening linkers contained no more than 12 residues. Candidate windows of 8–65 residues were generated by extending up to six residues beyond the contact block, using pseudorandom increments of one to three residues and a base seed of 20260526.

Candidates were first retained using one of three geometric criteria. Core candidates required local contact fraction, contact-block coverage and global contact coverage of at least 0.60, 0.45 and 0.10, respectively, and a non-contact fraction no greater than 0.45. Helical candidates—defined by at least 50% of (i,i + 4) Cα distances being no greater than 6.5 Å—used corresponding thresholds of 0.25, 0.35, 0.10 and 0.70. Bridge candidates were required to span at least two contact segments and used corresponding thresholds of 0.35, 0.50, 0.15 and 0.65.

Geometrically accepted candidates were subjected to four successive physical filters. First, hydrophobic binding-site residues were connected using a 9.5-Å Cα cutoff, and each of the two largest components was required to contain at least three residues. Second, the positive and negative intracomponent frustration sums of the largest component were normalized by its residue count. Candidates were retained when the normalized negative score was between −0.4 and 0, the normalized positive score was between 0 and 3, and the total positive score was between 0 and 75.

Third, the component trunk was defined as the shortest path connecting the most distant endpoints of its Cα graph. Positive configurational-frustration indices were summed for each trunk residue. Candidates were required to have a trunk of 3–10 residues and a mean residue-level positive-frustration sum below 15.

Finally, the largest hydrophobic component was designated S0, and non-overlapping shells S1 and S2 were constructed using a 4.5-Å heavy-atom cutoff. Each shell score was the mean residue-level positive-frustration sum. Candidates were retained when the score difference from S0 to S1 was greater than −6.5 and that from S1 to S2 was less than 1.5, yielding 15,325 protein-derived peptide–receptor complexes.

### Ourotide structure generation and affinity prediction

#### Peptide structure generation

Ourotide represents receptor–peptide complexes at residue-level Cα resolution. Node features combine amino-acid identity, local backbone geometry and residue embeddings generated by ESM-2 (esm2_t12_35M_UR50D).^50^ The receptor ESM-2 representation was kept frozen, whereas the final two transformer layers used to encode peptide sequences were fine-tuned during training. Graph edges encode Cα distances using radial basis functions together with global and local directions, relative orientations between residue frames, residue-pair identities, sequence separation, chain adjacency and chain identity. Receptor representations were processed by four graph-neural-network layers, and the conditional vector field was parameterized by six layers that exchanged information within the peptide and across the receptor–peptide interface.

Structure generation was conditioned on the receptor structure, receptor sequence, peptide sequence and an optional receptor hotspot. An anchor module assigned a probability to each receptor residue and predicted an initial orientation for the peptide. A Cα chain with an approximately 3.8-Å spacing was initialized around the predicted anchor and perturbed with Gaussian noise with a standard deviation of 5 Å. During training, hotspot annotations derived from the corresponding training complexes were provided to the model. During inference, hotspots could be specified by the user or predicted from the receptor using the hydrophobic-connectivity and configurational-frustration procedure described above. When no binding region was specified, an all-zero hotspot mask and the receptor Cα centroid were used, enabling inference without site-specific input.

The structure generator was trained by conditional flow matching^51^ along a polynomial interpolation between the initialized coordinates and the target peptide coordinates in each training complex. Self-conditioning was applied to 50% of the training samples. The training objective combined flow-matching, coordinate-reconstruction, frame-aligned point-error, peptide-chain geometry, steric-clash, receptor–peptide distance and anchor-prediction losses. Their respective weights were 1.0, 0.5, 1.0, 0.2, 0.2, 10.0 and 0.1. The receptor–peptide distance loss considered target distances below 12 Å and retained at most 60 receptor neighbours for each peptide residue. The physics-guided dataset was divided into 12,771 training, 1,277 validation and 1,277 held-out test complexes. The model was optimized using AdamW^52^ with a batch size of 16, an initial learning rate of 2 × 10−4, a weight decay of 10−4, gradient clipping at 1.0 and a cosine learning-rate schedule following 1,000 warm-up steps. The checkpoint with the lowest validation peptide Cα-RMSD was retained. At inference, the learned vector field was integrated using 20 Heun predictor–corrector steps to generate peptide conformations.

#### Binding-affinity prediction

The affinity model represents each receptor–peptide complex as a Cα-level geometric graph. Residue features comprise amino-acid identity, chain identity and frozen ESM-2 embeddings. Separate receptor–receptor, peptide– peptide and bidirectional receptor–peptide edges were constructed from frame-aware geometric features. Four message-passing layers were used to update the receptor and peptide representations. The final complex representation combined attention pooling over receptor–peptide edges, gated pooling of receptor and peptide residues, and seven interface-level geometric descriptors encoding receptor–peptide contacts, close contacts, steric clashes, hydrophobic contacts, the mean residue-wise minimum distance, the global minimum interchain distance and relative chain size. Separate regression heads predicted pKd, Rosetta energy and AWSEM energy.

Affinity training was performed in two stages. First, the model was pretrained on the physics-guided augmented complexes using their normalized Rosetta and AWSEM energies as supervision.^27,28^ Rosetta and AWSEM losses were assigned weights of 1.0 and 0.25, respectively. The pretrained model was subsequently fine-tuned on experimental receptor–peptide complexes with measured Kd, converted to pKd = −log10(Kd [M]). During fine-tuning, an energy-labelled batch from the augmented dataset was introduced every three optimization steps to retain the energetic representations learned during pretraining. The pKd, Rosetta and AWSEM loss weights during this stage were 1.0, 0.05 and 0.02, respectively, and all three regression tasks used the Huber loss.

Experimental-affinity complexes were grouped according to receptor-sequence similarity and divided into five folds. After removing close neighbours of the external test set, 213 experimental Kd complexes distributed across 124 receptor clusters were retained for model development. Each fold was trained with two random seeds, producing ten models in total. Their mean prediction was used as the final pKd, and the standard deviation among models was used to quantify ensemble uncertainty.

### Benchmarking and evaluation metrics

#### Structure-prediction benchmark

A benchmark of 200 receptor–peptide complexes comprising 200 distinct receptor entries was selected from the held-out structure test set, with peptide lengths ranging from 8 to 65 residues. Ourotide was compared with DynamicBind, PepFlow, PepGLAD, AfDesign, RFdiffusion and AlphaFold 3 using the input protocols supported by each method.^11,21–26^ Ourotide, AfDesign and RFdiffusion were conditioned on receptor hotspot residues; PepFlow used receptor residues surrounding the hotspot region; PepGLAD used a receptor pocket defined from the reference binding region; and DynamicBind and AlphaFold 3 received the complete receptor. The native peptide sequence was supplied to sequence-conditioned structure-prediction methods, whereas peptide length was specified for backbone-generative methods. For the primary structural comparison, five candidate conformations were evaluated per target for each method. Comparisons were performed on strict common sets for which all included methods produced valid structures. The seven-method comparison comprised 47 complexes within the shared peptide-length range, whereas the five-method comparison involving Ourotide, DynamicBind, AfDesign, RFdiffusion and AlphaFold 3 comprised 138 complexes across the full length range. Method coverage and the numbers of valid candidates are reported in the Supplementary Information.

#### Structure evaluation

Predicted complexes were first superposed on their reference structures by least-squares alignment of matched receptor Cα atoms using the Kabsch algorithm.^53^ Peptide Cα root-mean-square deviation (Cα-RMSD) was then calculated without independently fitting the peptide, with peptide residues matched in sequence order. For each receptor–peptide complex, the lowest Cα-RMSD among the five candidate conformations was retained as the best-of-five structural result. Interface recovery was evaluated using Q_interface. Native receptor–peptide contacts were defined as residue pairs with a Cα–Cα distance of no more than 9.5 Å. For each native contact, a Gaussian distance-similarity term was calculated from the difference between its predicted and reference distances, with the distance tolerance scaled as 1.0 Å × (|i − j| + 1)^0.15. (Q_{\mathrm{interface}}) was calculated as the mean similarity over all native interchain contacts and ranged from 0 to 1, with higher values indicating more accurate recovery of the reference interface. The strict common Q_interface sets contained 31 complexes for the seven-method comparison and 138 complexes for the five-method comparison.

#### Binding-affinity benchmark

Binding-affinity prediction was evaluated on a locked test set of 49 receptor–peptide complexes with experimentally determined Kd values. Measurements were converted to molar units and transformed as pKd = −log10(Kd [M]). The final Ourotide prediction was obtained by averaging the outputs of ten models trained across five receptor-clustered folds and two random seeds. Ourotide was compared with Rosetta, AWSEM, GROMACS, DynamicBind, PeptiVerse and PepPAP on the same 49 complexes.^21,27–31^ PeptiVerse and PepPAP predictions were obtained using their published models and corresponding inference procedures. Affinity accuracy was evaluated using the absolute error |ΔpKd|, with success thresholds of 0.5, 1.0 and 2.0 pKd units. Success rates were calculated using all 49 complexes as the denominator, with unavailable predictions counted as failures.

## Data availability

The experimentally determined structures analysed in this study are available from the Protein Data Bank. The accession codes, raw input files, processed datasets, benchmark predictions and source data underlying the figures will be made publicly available upon publication.

## Code availability

The source code used for physics-guided data construction, Ourotide model training and inference, benchmark evaluation and data analysis will be made publicly available upon publication at https://github.com/yuhangshen2001/ourotide and Zenodo.

## Funding

This work was supported by the Innovative Drug Research and Development programme of the National Science and Technology Major Project (grant no. 2025ZD1801601 to X.C.), Nanjing Medical University (grant no. NMUR 2024007 to X.C.), and the Natural Science Foundation of Jiangsu Province (grant no. BK20211254 to N.J.).

## Author contributions

Y.S. conceived the study, developed the Ourotide framework, curated the datasets, performed the computational analyses and molecular simulations, evaluated the models, prepared the figures and wrote the original manuscript. J.Z. contributed to method development, software implementation, computational experiments and validation. Z.W. contributed to data curation, model evaluation and result validation. Z.X. contributed to dataset construction and computational analysis. Q.Y. contributed to data processing and benchmark evaluation. W.Z. contributed to data collection and result validation. Q.Z. and F.H. provided scientific guidance, resources and supervision, and reviewed the manuscript. N.J. contributed to study conceptualization, funding acquisition, project administration and supervision, and revised the manuscript. X.C. conceived and directed the study, supervised method development and data interpretation, acquired funding, and revised the manuscript. All authors discussed the results and approved the final manuscript.

## Competing interests

The authors declare no competing interests.

## Additional information

Supplementary information is available for this paper. Correspondence should be addressed to Qigang Zhou, Feng Han, Nan Jiang or Xun Chen. Requests for data, code and materials should be addressed to Yuhang Shen.

## Notes

### Competing Interest Statement

The authors have declared no competing interest.

## References

1. Muttenthaler, M., King, G. F., Adams, D. J. & Alewood, P. F. Trends in peptide drug discovery. Nat. Rev. Drug Discov. 20, 309–325 (2021).

2. London, N., Movshovitz-Attias, D. & Schueler-Furman, O. The structural basis of peptide–protein binding strategies. Structure 18, 188–199 (2010).

3. Xu, S. et al. Benchmarking all-atom biomolecular structure prediction with FoldBench. Nat. Commun. 17, 442 (2026).

4. Masters, M. R., Mahmoud, A. H. & Lill, M. A. Investigating whether deep learning models for co-folding learn the physics of protein–ligand interactions. Nat. Commun. 16, 8854 (2025).

5. Lau, J. L. & Dunn, M. K. Therapeutic peptides: historical perspectives, current development trends, and future directions. Bioorg. Med. Chem. 26, 2700–2707 (2018).

6. Wang, L. et al. Therapeutic peptides: current applications and future directions. Signal Transduct. Target. Ther. 7, 48 (2022).

7. Ciemny, M., et al. Protein–peptide docking: opportunities and challenges. Drug Discov. Today 23, 1530–1537 (2018).

8. Wright, P. E. & Dyson, H. J. Linking folding and binding. Curr. Opin. Struct. Biol. 19, 31–38 (2009).

9. Mobley, D. L. & Gilson, M. K. Predicting binding free energies: frontiers and benchmarks. Annu. Rev. Biophys. 46, 531–558 (2017).

10. Genheden, S. & Ryde, U. The MM/PBSA and MM/GBSA methods to estimate ligand-binding affinities. Expert Opin. Drug Discov. 10, 449–461 (2015).

11. Abramson, J. et al. Accurate structure prediction of biomolecular interactions with AlphaFold 3. Nature 630, 493–500 (2024).

12. Krishna, R. et al. Generalized biomolecular modeling and design with RoseTTAFold All-Atom. Science 384, eadl2528 (2024).

13. Joeres, R., Blumenthal, D. B. & Kalinina, O. V. Data splitting to avoid information leakage with DataSAIL. Nat. Commun. 16, 3337 (2025).

14. Walsh, I. et al. DOME: recommendations for supervised machine learning validation in biology. Nat. Methods 18, 1122–1127 (2021).

15. Tsaban, T. et al. Harnessing protein folding neural networks for peptide–protein docking. Nat. Commun. 13, 176 (2022).

16. Weng, G. et al. Comprehensive evaluation of fourteen docking programs on protein–peptide complexes. J. Chem. Theory Comput. 16, 3959–3969 (2020).

17. Wen, Z., He, J., Tao, H. & Huang, S.-Y. PepBDB: a comprehensive structural database of biological peptide–protein interactions. Bioinformatics 35, 175–177 (2019).

18. Martins, P. M. et al. Propedia: a database for protein–peptide identification based on a hybrid clustering algorithm. BMC Bioinformatics 22, 1 (2021).

19. Zhu, N., et al. A data resource of peptide–protein interaction structures and binding affinities for peptide drug discovery. Sci. Data 10.1038/s41597-026-07962-1 (2026).

20. Wang, S. et al. A structure-based data set of protein–peptide affinities and its nonredundant benchmark: potential applications in computational peptidology. Curr. Med. Chem. 31, 4127–4137 (2024).

21. Lu, W. et al. DynamicBind: predicting ligand-specific protein–ligand complex structure with a deep equivariant generative model. Nat. Commun. 15, 1071 (2024).

22. Li, J. et al. Full-atom peptide design based on multi-modal flow matching. In Proc. 41st Int. Conf. Mach. Learn. 235, 27615–27640 (PMLR, 2024).

23. Kong, X., Jia, Y., Huang, W. & Liu, Y. Full-atom peptide design with geometric latent diffusion. Adv. Neural Inf. Process. Syst. 37, 74808–74839 (2024).

24. Wang, J. et al. Scaffolding protein functional sites using deep learning. Science 377, 387–394 (2022).

25. Ovchinnikov, S. & Dauparas, J. AfDesign—partial hallucination with sidechain constraints. Zenodo 10.5281/zenodo.6803187 (2022).

26. Watson, J. L. et al. De novo design of protein structure and function with RFdiffusion. Nature 620, 1089–1100 (2023).

27. Alford, R. F. et al. The Rosetta all-atom energy function for macromolecular modeling and design. J. Chem. Theory Comput. 13, 3031–3048 (2017).

28. Davtyan, A. et al. AWSEM-MD: protein structure prediction using coarse-grained physical potentials and bioinformatically based local structure biasing. J. Phys. Chem. B 116, 8494–8503 (2012).

29. Abraham, M. J. et al. GROMACS: high performance molecular simulations through multi-level parallelism from laptops to supercomputers. SoftwareX 1–2, 19–25 (2015).

30. Zhang, Y. et al. PeptiVerse: a unified platform for therapeutic peptide property prediction. Nat. Commun. 17, 6819 (2026).

31. Sun, X., Wu, Z., Su, J. & Li, C. A deep attention model for wide-genome protein– peptide binding affinity prediction at a sequence level. Int. J. Biol. Macromol. 276, 133811 (2024).

32. Clackson, T. & Wells, J. A. A hot spot of binding energy in a hormone–receptor interface. Science 267, 383–386 (1995).

33. Chandler, D. Interfaces and the driving force of hydrophobic assembly. Nature 437, 640–647 (2005).

34. Levy, Y. & Onuchic, J. N. Water mediation in protein folding and molecular recognition. Annu. Rev. Biophys. Biomol. Struct. 35, 389–415 (2006).

35. Hoffmann, J. et al. Training compute-optimal large language models. Adv. Neural Inf. Process. Syst. 35, 30016–30030 (2022).

36. Wallach, I. & Heifets, A. Most ligand-based classification benchmarks reward memorization rather than generalization. J. Chem. Inf. Model. 58, 916–932 (2018).

37. Wang, H. et al. Scientific discovery in the age of artificial intelligence. Nature 620, 47–60 (2023).

## Methods references

38. Wang, R., Fang, X., Lu, Y. & Wang, S. The PDBbind database: collection of binding affinities for protein–ligand complexes with known three-dimensional structures. J. Med. Chem. 47, 2977–2980 (2004).

39. Parra, R. G. et al. Protein Frustratometer 2: a tool to localize energetic frustration in protein molecules, now with electrostatics. Nucleic Acids Res. 44, W356–W360 (2016).

40. Lee, J. et al. CHARMM-GUI Input Generator for NAMD, GROMACS, AMBER, OpenMM, and CHARMM/OpenMM simulations using the CHARMM36 additive force field. J. Chem. Theory Comput. 12, 405–413 (2016).

41. Jorgensen, W. L., Chandrasekhar, J., Madura, J. D., Impey, R. W. & Klein, M. L. Comparison of simple potential functions for simulating liquid water. J. Chem. Phys. 79, 926–935 (1983).

42. Huang, J. et al. CHARMM36m: an improved force field for folded and intrinsically disordered proteins. Nat. Methods 14, 71–73 (2017).

43. Vanommeslaeghe, K. et al. CHARMM general force field: a force field for drug-like molecules compatible with the CHARMM all-atom additive biological force fields. J. Comput. Chem. 31, 671–690 (2010).

44. Bussi, G., Donadio, D. & Parrinello, M. Canonical sampling through velocity rescaling. J. Chem. Phys. 126, 014101 (2007).

45. Bernetti, M. & Bussi, G. Pressure control using stochastic cell rescaling. J. Chem. Phys. 153, 114107 (2020).

46. Essmann, U. et al. A smooth particle mesh Ewald method. J. Chem. Phys. 103, 8577–8593 (1995).

47. Hess, B., Bekker, H., Berendsen, H. J. C. & Fraaije, J. G. E. M. LINCS: a linear constraint solver for molecular simulations. J. Comput. Chem. 18, 1463–1472 (1997).

48. Lu, W. et al. OpenAWSEM with Open3SPN2: a fast, flexible, and accessible framework for large-scale coarse-grained biomolecular simulations. PLoS Comput. Biol. 17, e1008308 (2021).

49. Kumar, S., Rosenberg, J. M., Bouzida, D., Swendsen, R. H. & Kollman, P. A. The weighted histogram analysis method for free-energy calculations on biomolecules. I. The method. J. Comput. Chem. 13, 1011–1021 (1992).

50. Lin, Z. et al. Evolutionary-scale prediction of atomic-level protein structure with a language model. Science 379, 1123–1130 (2023).

51. Lipman, Y., Chen, R. T. Q., Ben-Hamu, H., Nickel, M. & Le, M. Flow matching for generative modeling. In International Conference on Learning Representations https://openreview.net/forum?id=PqvMRDCJT9t (2023).

52. Loshchilov, I. & Hutter, F. Decoupled weight decay regularization. In International Conference on Learning Representations https://openreview.net/forum?id=Bkg6RiCqY7 (2019).

53. Kabsch, W. A solution for the best rotation to relate two sets of vectors. Acta Crystallogr. A 32, 922–923 (1976).

