## Supplementary Information for "Ourotide: decoding the hierarchical peptide recognition for generative design"

#### Contents


- Supplementary Methods
- Supplementary Figures 1–28
- Supplementary Tables
- Supplementary References

#### Supplementary Methods

##### 1. Construction of the reference complex libraries

###### Protein–peptide and protein–protein reference libraries

Protein–peptide and protein–protein complexes were collected from the protein–protein subset of PDBbind 2020 using the INDEX_general_PP.2020 index and the corresponding coordinate files. Records reporting Kd, Ki or IC50 values were converted to molar units. An operational affinity index was calculated as −log10(X [M]), where X denotes the reported Kd, Ki or IC50 value, and entries with an affinity index greater than 7 were retained. For measurements reported as inequalities, the threshold was applied to the numerical bound recorded in the PDBbind index, and the original measurement qualifier was retained in the dataset metadata.[38,54]

Peptidic chains were identified from residues containing the backbone atoms N, Cα, C and O. Structures were required to contain exactly two peptidic chains composed of the 20 standard amino acids. The shorter chain was designated as the ligand and the longer chain as the receptor; when the two chains had the same length, chain identifiers were used to obtain a deterministic assignment. Complexes in which the ligand chain contained 8–65 residues were assigned to the protein–peptide reference library, yielding 186 complexes. Complexes in which the shorter chain contained more than 65 residues were assigned to the protein–protein reference library, yielding 246 complexes. FDA approval status was not used to construct either library.

###### Protein–small-molecule reference library

Candidate protein–small-molecule complexes were collected from the PDBbind 2020 refined set and the remaining protein–ligand entries in the PDBbind 2020 general set. These source collections contained 5,317 and 14,128 entries, respectively. Each entry was required to contain a protein coordinate file and a ligand structure file. Protein structures were retained when exactly one peptidic chain was detected and the detected chain contained only standard amino-acid residues. Entries containing more than one molecular record in the ligand SDF file were excluded. This structural eligibility screening retained 3,055 complexes from the refined set and 8,287 complexes from the remaining protein–ligand set, producing 11,342 structurally eligible candidate complexes.[38]

An in-house approved-drug table was filtered to retain records whose approval-status field contained 'FDA'. SMILES strings for the retained drugs were processed with RDKit by removing explicit hydrogen atoms and generating canonical isomeric SMILES. For each PDBbind candidate, the corresponding ligand was read from its SDF file, with a MOL2 representation used when required. Ligands were sanitized where possible, explicit hydrogen atoms were removed, and canonical isomeric SMILES were generated using the same RDKit procedure. PDBbind ligands were matched to the FDA-approved drug table by exact equality of canonical isomeric SMILES. Multiple database matches corresponding to the same PDB entry were collapsed to a single complex.[55]

Experimental Kd, Ki or IC50 annotations for the matched complexes were retrieved from INDEX_general_PL_data.2020. Entries with a PDBbind affinity index greater than 7 were retained. The intersection of the structural eligibility, FDA-approved-drug matching and affinity-index criteria yielded 387 protein–small-molecule complexes, which constituted the small-molecule reference library used for comparison with the protein–peptide and protein–protein binding sites.

**Affinity-unrestricted control libraries for Supplementary analyses**

To evaluate whether the structural differences among peptide-binding sites, small-molecule pockets and protein–protein interfaces depended on the affinity-index threshold used for the primary reference libraries, we constructed three additional affinity-unrestricted control libraries. These control libraries were assembled without requiring a reported Kd, Ki or IC50, without applying an affinity-index threshold and, for the small-molecule library, without restricting ligands to FDA-approved drugs. Affinity annotations were retained as metadata when available but were not used to select these complexes.

The affinity-unrestricted protein–peptide and protein–protein control libraries were constructed from the protein–protein subset of PDBbind 2020 using INDEX_general_PP.2020 and the corresponding coordinate files. Entries were required to contain exactly two peptidic chains. Chain lengths were calculated from residues belonging to the 20 standard amino acids and containing the backbone atoms N, Cα, C and O, and complexes containing detected non-standard peptidic residues were excluded. The shorter chain was assigned as the ligand and the longer chain as the receptor; ties were resolved deterministically according to chain identifier. Complexes with a shorter chain containing fewer than eight residues were excluded. Those with a shorter chain of 8–65 residues were assigned to the affinity-unrestricted protein–peptide control library (n = 349), whereas those with a shorter chain longer than 65 residues were assigned to the affinity-unrestricted protein–protein control library (n = 483). For subsequent calculations, receptor and ligand chains were renamed A and Z, respectively, and retained residues were renumbered consecutively from residue 1.[38]

The affinity-unrestricted protein–small-molecule control library was constructed from the PDBbind 2020 refined set, which contained 5,317 source entries. Entries were required to contain a protein coordinate file and a ligand SDF file. Protein structures were retained when exactly one peptidic chain was detected and the detected chain contained only standard amino-acid residues with complete N, Cα, C and O backbone atoms. Ligand files were required to contain a single molecular record. This screening retained 3,055 protein–small-molecule complexes. For downstream analysis, the receptor chain was renamed A. Ligand coordinates were read from the corresponding SDF file and written as a single residue named UNL in chain Z with residue number 999. The standardized receptor and ligand coordinates were then combined into a common complex representation.[38]

These affinity-unrestricted libraries were used exclusively in the Supplementary comparative analyses to assess the robustness of the observed structural and energetic trends to affinity-based dataset selection. They were not used to construct the primary high-affinity-index reference libraries or to train Ourotide.

#### 2. Binding-site geometry and recognition-regime assignment

##### Binding-site identification and residue classification

All reference complexes were converted to a common chain convention in which the receptor was represented by chain A and the peptide, protein or small-molecule ligand by chain Z. A receptor residue was assigned to the binding site when the minimum distance between any atom of that residue and any ligand atom was no greater than 6.5 Å. Ala, Val, Leu, Ile, Met, Phe, Trp and Pro were classified as hydrophobic; Asp, Glu, Lys, Arg, Asn, Gln, Ser and Thr were classified as hydrophilic; and Gly, Cys, Tyr and His were classified as neutral. Residue identities, sequence numbers and physicochemical classes were retained for subsequent geometric and connectivity analyses.

##### Binding-site connectivity

Binding-site residues were represented as graph nodes. Graphs were constructed separately for hydrophobic and hydrophilic residues, with an edge placed between two nodes when the distance between their Cα atoms was no greater than 9.5 Å. Connected components were identified from each graph and ranked by their number of residues. The number and size of hydrophobic connected components were used to quantify hydrophobic-site multiplicity. Unless otherwise indicated, the component-based analyses and recognition-regime assignments reported in this study refer to the hydrophobic graph.

##### Binding-site geometry

Binding-site geometry was calculated from the Cα coordinates of all receptor residues assigned to the 6.5-Å binding site. The coordinates were centred, and principal-component analysis was used to obtain three mutually orthogonal axes with covariance eigenvalues $\lambda 1 \geq \lambda 2 \geq \lambda 3$. The binding-site flatness index was defined as $F = 1 - \lambda 3/\lambda 1$, such that larger values indicate a more planar distribution of binding-site residues. Binding-site thickness was defined as $T = \sqrt{\lambda 3}$ and was reported in ångströms. The binding-site hydrophobic fraction was calculated as the number of hydrophobic binding-site residues divided by the total number of binding-site residues. Numerical array operations and eigendecomposition were performed using NumPy.[56]

##### Peptide engagement and recognition-regime assignment

For protein–peptide complexes, peptide engagement was defined as the fraction of peptide residues containing at least one atom within 6.5 Å of any receptor atom. The joint distribution of peptide engagement and hydrophobic-component number was used to define the core regions of three operational recognition regimes. Complexes with peptide engagement below 0.5 were assigned to the protein-like regime (regime iii). Among complexes with peptide engagement between 0.7 and 1.0, those containing one hydrophobic connected component were assigned to the small-molecule-like regime (regime ii), whereas those containing at least two hydrophobic connected components were assigned to the peptide-specific regime characterized by a hierarchical subpocket architecture (regime i). Complexes outside these core regions were excluded from analyses requiring an unambiguous regime assignment but were retained in analyses of the overall peptide-site distribution where indicated.

##### 3. All-atom molecular-dynamics simulations

All-atom molecular-dynamics simulations were performed for representative small-molecule, peptide and protein complexes 1MUI, 1DPJ and 1A22, respectively, using GROMACS 2024.5. The systems were prepared with CHARMM-GUI 3.7 in periodic cubic boxes containing CHARMM-modified TIP3P water and 0.15 M NaCl. Proteins and peptides were described with the CHARMM36m force field, and the small-molecule ligand in 1MUI was parameterized with the CHARMM General Force Field.[29,40–43]

Each system was subjected to position-restrained steepest-descent energy minimization for at most 50,000 steps, using a maximum-force tolerance of 100 kJ mol−1 nm−1. Protein and peptide backbone and side-chain heavy atoms were restrained with force constants of 400 and 40 kJ mol−1 nm−2, respectively, and small-molecule heavy atoms were restrained with a force constant of 400 kJ mol−1 nm−2. The minimized systems were equilibrated for 125 ps in the NVT ensemble using a 1-fs integration timestep. Temperature was maintained at 300 K using the velocity-rescaling thermostat, with solute and solvent coupled separately and a coupling time constant of 1.0 ps. Initial velocities were generated independently for the five replicas using random seeds 1001, 1002, 1003, 1004 and 1005.[44]

Unrestrained production simulations were performed for 500 ns in the NPT ensemble using a 2-fs timestep. Temperature was maintained at 300 K using velocity rescaling with a coupling time constant of 1.0 ps. Pressure was maintained isotropically at 1 bar using the stochastic cell-rescaling barostat, with a coupling time constant of 5.0 ps and a compressibility of 4.5 × 10−5 bar−1. Long-range electrostatic interactions were treated with particle-mesh Ewald summation using a 1.2-nm real-space cutoff. Van der Waals interactions were force-switched between 1.0 and 1.2 nm, and the neighbour-list cutoff was 1.2 nm. Bonds involving hydrogen atoms were constrained using LINCS.

Centre-of-mass motion was removed separately for the solute and solvent groups every 100 steps. Compressed coordinates were recorded every 50 ps.[44–47]

Five independent 500-ns production trajectories were generated for each complex, corresponding to an aggregate sampling time of 2.5 μs per system. Unless otherwise indicated, structural, interfacial and hydration analyses were performed using the final 200 ns of each trajectory.

**Trajectory-derived interfacial geometry, hydration and mechanical metrics**

The same trajectory-derived metrics were calculated for two complementary groupings. Study A compared peptide-specific sites, small-molecule sites and protein interfaces, whereas study B compared the largest, second-largest and third-largest hydrophobic components within peptide-specific sites.

Buried surface area (BSA) was calculated for each frame as BSA = (SASA_pocket + SASA_binder − SASA_pocket+binder)/2, where the three solvent-accessible surface areas were obtained from GROMACS analyses of the pocket, binder and combined pocket–binder groups, respectively. BSA was reported in $nm^2$. To quantify relative changes within each system, the BSA ratio was calculated by dividing the value in each frame by that in the first analysed frame.

Shared interfacial waters were defined as water oxygen atoms located within 0.35 nm of both pocket heavy atoms and binder heavy atoms in the same frame. A shared-water density proxy was calculated as the number of shared waters divided by the convex-hull volume of their oxygen coordinates and was expressed in $nm^{-3}$. Positive density values were transformed to log10 scale for visualization.

For force analysis, the instantaneous net force on the pocket was obtained by vectorially summing the forces over all pocket atoms. The magnitude of this vector was evaluated in each frame, and the pocket net-force root-mean-square value was calculated over consecutive non-overlapping windows of 500 frames. Values were reported in kJ $mol^{-1}$ $nm^{-1}$; thus, each 5,000-frame trajectory segment contributed ten window-level observations.

Local mechanical stress was estimated from the pocket heavy atoms. Their convex-hull volume V and centre-of-mass-referenced coordinates were used to construct the configurational stress tensor $\sigma = -V^{-1} \Sigma i[(ri - rCOM) \otimes Fi]$. Tensor elements were converted from kJ $mol^{-1}$ $nm^{-3}$ to bar using a factor of 16.60539067. The local hydrostatic pressure was calculated as $p = -Tr(\sigma)/3$. The deviatoric tensor was defined as $s = \sigma - Tr(\sigma)I/3$, and the von Mises stress was calculated as $[3(s:s)/2]^{1/2}$; both quantities were reported in bar. Stress anisotropy was calculated from the ordered eigenvalues $\lambda 1 \geq \lambda 2 \geq \lambda 3$ of σ as $(\lambda 1 - \lambda 3)/(|\lambda 1| + |\lambda 2| + |\lambda 3| + \varepsilon)$, with $\varepsilon = 10^{-12}$. Stress anisotropy is dimensionless. Convex-hull calculations were performed using SciPy.[57]

#### 4. Physics-guided peptide data augmentation

##### Candidate extraction from protein–protein complexes

Candidate peptide–receptor complexes were extracted from the source protein–protein library described above. For each complex containing two eligible protein chains, both chain orientations were considered, with the putative receptor required to contain at least 66 standard amino-acid residues. A residue on the putative ligand chain was classified as interfacial when it formed at least three interchain heavy-atom pairs within 4.5 Å of the receptor. Coordinate parsing and heavy-atom neighbour searches were performed using Biopython.[58]

Interfacial residues were partitioned into contact segments according to their positions along the ligand chain. Contact residues separated by no more than three sequence positions were assigned to the same segment. Consecutive blocks containing up to three contact segments were enumerated, provided that adjacent segments were separated by no more than 12 residues and that the complete block contained at least three interfacial residues. Candidate windows of 8–65 residues were sampled within six residues on either side of each block. Windows containing discontinuities greater than two residue numbers were excluded.

Candidate windows were retained when they satisfied at least one of three geometric criteria. Core candidates required a contact density of at least 0.60, coverage of at least 0.45 of the contacts in the corresponding block, coverage of at least 0.10 of all ligand-chain contacts and a non-contact fraction no greater than 0.45. Helical candidates were identified when at least 50% of available $C\alpha i$–$C\alpha i+4$ distances were no greater than 6.5 Å; these candidates additionally required a contact density of at least 0.25, block-contact coverage of at least 0.35, global-contact coverage of at least 0.10 and a non-contact fraction no greater than 0.70. Bridge candidates were required to span at least two contact segments and to have a contact density of at least 0.35, block-contact coverage of at least 0.50, global-contact coverage of at least 0.15 and a non-contact fraction no greater than 0.65. Candidate start positions and lengths were sampled using deterministic increments of one to three residues with a base random seed of 20260526. Duplicate windows with identical start and end positions were collapsed by retaining the assignment with the highest block coverage, contact density and contact count.

**Hydrophobic-component screening**

For each geometrically accepted candidate, receptor residues containing at least one heavy atom within 6.5 Å of a candidate-peptide heavy atom were assigned to the binding site. Ala, Val, Leu, Ile, Met, Phe, Trp and Pro were classified as hydrophobic. Hydrophobic binding-site residues were represented as graph nodes and connected when their Cα atoms were within 9.5 Å. Connected components were ranked by residue count, and candidates were retained when each of the two largest components contained at least three residues.

**Component-level energetic screening**

A precomputed pairwise configurational-frustration matrix was used to characterize the largest hydrophobic component. For a component containing nC residues, positive and negative pairwise frustration values in the upper triangle of the component submatrix were summed separately. Component-level averages were calculated by dividing each sum by nC. Candidates were retained when the average negative score was between −0.4 and 0, the average positive score was between 0 and 3, and the positive-score sum was between 0 and 75.[28,39]

**Component-trunk screening**

The largest hydrophobic component was represented using the same 9.5-Å Cα graph. Terminal nodes were defined as degree-one nodes; when fewer than two such nodes were present, nodes with maximum graph eccentricity were used. The most distant endpoint pairs were identified by graph distance, and shortest paths connecting these endpoints were enumerated. For each component residue, a residue-level frustration score was calculated by summing its non-zero pairwise frustration values across the precomputed receptor matrix. When multiple equivalent shortest paths were present, a deterministic ranking based on endpoint and internal-residue frustration scores was used. The selected trunk was required to contain 3–10 residues and to have a mean residue-level frustration score below 15. Graph construction and shortest-path analysis were performed using NetworkX.[59]

**Shell-level energetic screening**

The largest hydrophobic component was designated shell S0. Successive, non-overlapping receptor shells were generated iteratively by assigning residues containing a heavy atom within 4.5 Å of the preceding shell. For each shell, the mean residue-level frustration score was calculated over its constituent residues. Candidates were retained when the S1-minus-S0 score difference was greater than −6.5 and the S2-minus-S1 difference was less than 1.5.

**Binding-site flatness screening**

After completion of the component- and shell-level filters, binding-site flatness was evaluated from the Cα coordinates of the receptor residues within 6.5 Å of the candidate peptide. The coordinates were centred and subjected to principal-component analysis, yielding covariance eigenvalues $\lambda 1 \geq \lambda 2 \geq \lambda 3$. The flatness index was calculated as $F = 1 - \lambda 3/\lambda 1$, and candidates with F greater than 0.8 were retained. Application of the contact-geometry, hydrophobic-component, component-energy, trunk, shell and final flatness criteria yielded 15,325 physics-guided peptide–receptor complexes.

**Geometry-derived augmentation control**

A geometry-derived augmentation control was constructed from the candidate windows obtained after the contact-based geometric extraction stage and before application of the hydrophobic-component, component-energy, trunk, shell and flatness acceptance criteria. Candidates were processed separately for each source complex and chain orientation. They were ranked by decreasing interfacial-contact count, local contact ratio and largest-component size, followed by increasing average negative-frustration score and candidate identifier. For each peptide length, the highest-ranked candidate was retained.

A maximum of ten candidates was selected for each chain orientation. When no more than ten unique peptide lengths were available, all length-unique candidates were retained. Otherwise, the observed peptide-length range was divided into ten bins and the highest-ranked candidate from each occupied bin was selected. Unfilled positions were assigned iteratively to candidates that maximized the minimum length difference from the already selected set, with the same ranking criteria used to resolve ties. This control preserved the contact-based extraction and peptide-length diversity of the augmentation procedure without imposing the physical acceptance thresholds used to construct the physics-guided dataset.

#### 5. Ourotide structure generator

Ourotide generates receptor-bound peptide conformations using a residue-level, Cα-only conditional flow-matching model. Receptor and peptide residues were represented by their amino-acid identities, local Cα-chain geometry and 480-dimensional residue embeddings obtained from layer 6 of the pretrained ESM-2 model esm2_t12_35M_UR50D. The ESM-2 parameters were held fixed during structure-model training. All coordinates were centred using the receptor Cα centroid. During training, the receptor context was restricted to residues whose Cα atoms were within 30 Å of any peptide Cα atom, with maximum receptor and peptide lengths of 350 and 65 residues, respectively.[50]

The receptor and peptide were represented as frame-aware geometric graphs. Edge features encoded Cα distances using radial basis functions, global and local inter-residue directions, relative orientations between residue frames, amino-acid-pair identities, sequence separation, chain adjacency and chain identity. Receptor representations were processed by four message-passing layers. The conditional vector field comprised six successive blocks that exchanged information along peptide–peptide, peptide-to-receptor and receptor-to-peptide edges. The corresponding neighbourhood sizes were 6, 8 and 8 residues, respectively, whereas receptor–receptor graphs used 16 nearest neighbours. Node and edge representations had dimensions of 128 and 64, respectively, and a dropout probability of 0.1 was used.

Structure generation was conditioned on the receptor structure and sequence, the peptide sequence and a receptor hotspot. For supervised training and hotspot-conditioned benchmarking, receptor residues within 6.5 Å of the experimentally resolved peptide were first identified using heavy-atom distances. Ala, Val, Leu, Ile, Met, Phe, Trp and Pro within this interfacial region were represented as graph nodes and connected when their Cα atoms were within 9.5 Å; the largest connected hydrophobic component was used as the hotspot. The hotspot mask and its Cα centroid were supplied to an anchor module, which assigned a probability to each receptor residue and predicted an initial peptide orientation. The centre of the initial peptide distribution was calculated as the probability-weighted mean of the receptor Cα coordinates.

For receptor-only hotspot proposal, hydrophobic components containing at least three residues were enumerated over the receptor surface using the same 9.5-Å Cα connectivity criterion. Candidate components were ranked using a composite score integrating component-level configurational frustration, the length and frustration profile of the component trunk, energetic contrasts between successive 4.5-Å receptor shells and component size. The highest-ranked component and its Cα centroid were then used as the hotspot condition for peptide generation.

Initial peptide conformations were constructed as Cα chains with an adjacent-residue spacing of approximately 3.8 Å. Each chain was centred on the predicted anchor, oriented using the direction predicted by the anchor module and perturbed with isotropic Gaussian coordinate noise with a standard deviation of 5 Å. Conditional flow matching was performed along the polynomial path $x(t) = x(0) + [1 - (1 - t)^2][x(1) - x(0)]$, where $x(0)$ denotes the initialized peptide coordinates and $x(1)$ denotes the experimentally observed peptide coordinates. The corresponding target vector field was $2(1 - t)[x(1) - x(0)]$. Self-conditioning was applied to 50% of the training examples: a preliminary endpoint estimate was generated without gradient propagation and supplied as an additional geometric condition for the subsequent vector-field prediction.[51]

The training objective combined flow-matching, endpoint-coordinate reconstruction, Cα frame-aligned point-error, adjacent-residue chain-geometry, receptor–peptide steric-clash, receptor–peptide distance and anchor-prediction losses, with respective weights of 1.0, 0.5, 1.0, 0.2, 0.2, 10.0 and 0.1. The receptor–peptide distance loss considered native Cα pairs separated by less than 12 Å, retained at most 60 receptor neighbours for each peptide residue and weighted pairs according to $\exp(-d/4\ \text{Å})$. Anchor targets were receptor residues containing a Cα atom within 10 Å of any peptide Cα atom.

The physics-guided dataset comprised 12,771 training complexes, 1,277 validation complexes and 1,277 held-out test complexes. The model was optimized using AdamW on seven GPUs with a batch size of 16 per GPU, corresponding to an effective batch size of 112. The initial learning rate was $2 \times 10^{-4}$, with a weight decay of $10^{-4}$ and gradient clipping at 1.0. The learning rate was linearly warmed up over the first 1,000 optimization steps and subsequently decayed towards $10^{-5}$ using a cosine schedule. Training was performed using bfloat16 mixed precision with a random seed of 42. Model selection was based on peptide Cα-RMSD on the validation set. At inference, the learned vector field was integrated from $t = 0$ to $t = 1$ using 20 Heun predictor–corrector steps, and five independently initialized peptide conformations were evaluated for each target.[52]

#### 6. Ourotide affinity predictor

Ourotide represented each receptor–peptide complex as a residue-level Cα geometric graph. Node features comprised amino-acid identity, chain identity and frozen residue embeddings generated by the pretrained ESM-2 model esm2_t12_35M_UR50D. Separate receptor–receptor, peptide–peptide and bidirectional receptor–peptide edges were constructed from frame-aware geometric features. Receptor residues within 30 Å of the peptide were retained, with maximum receptor and peptide lengths of 350 and 65 residues, respectively. Four message-passing layers were used to update the receptor and peptide representations.[50]

The complex-level representation combined attention pooling over receptor–peptide edges, gated pooling over receptor and peptide residues, and seven interface descriptors: the fractions of receptor–peptide residue pairs within 8.0, 6.0 and 3.5 Å; the fraction of hydrophobic receptor–peptide contacts within 8.0 Å; the mean minimum receptor distance over peptide residues; the global minimum interchain distance; and the peptide-to-receptor length ratio. The resulting representation was passed to separate regression heads for pKd, Rosetta interface energy and AWSEM interaction energy.

Model training comprised energy-supervised pretraining followed by experimental-affinity fine-tuning. During pretraining, the physics-guided augmented complexes were labelled with calculated Rosetta and AWSEM energies. Rosetta interface energies were calculated using InterfaceAnalyzer with the ref2015 energy function, and dG_separated was used as the Rosetta target. AWSEM interaction energies were calculated as the energy of the complete complex minus the energies of the isolated receptor and peptide. Rosetta and AWSEM targets were normalized using statistics calculated from the corresponding training partition and optimized using Huber losses with weights of 1.0 and 0.25, respectively.[27,28]

The pretrained model was subsequently fine-tuned using receptor–peptide complexes with experimentally measured Kd values, converted to pKd = −log10(Kd [M]). During fine-tuning, one energy-labelled batch from the augmented dataset was included every three experimental-affinity optimization steps. The pKd, Rosetta and AWSEM loss weights were 1.0, 0.05 and 0.02, respectively, and all three regression tasks used the Huber loss. The encoder was optimized with a learning rate of $5 \times 10^{-5}$, whereas the pooling layers and regression heads used a learning rate of $3 \times 10^{-4}$. Models were selected according to validation pKd root-mean-square error.

Experimental-affinity complexes were clustered according to receptor-sequence similarity using a sequence-identity threshold of 40% and a minimum shorter-sequence coverage of 70%. After excluding close neighbours of the external test set, 213 complexes distributed across 124 receptor clusters were retained for model development and divided into five folds. Each fold was trained using two random seeds, producing ten models. The mean of their predictions was used as the final pKd estimate, and the standard deviation across the ensemble was used as an uncertainty estimate.

### Supplementary Figures

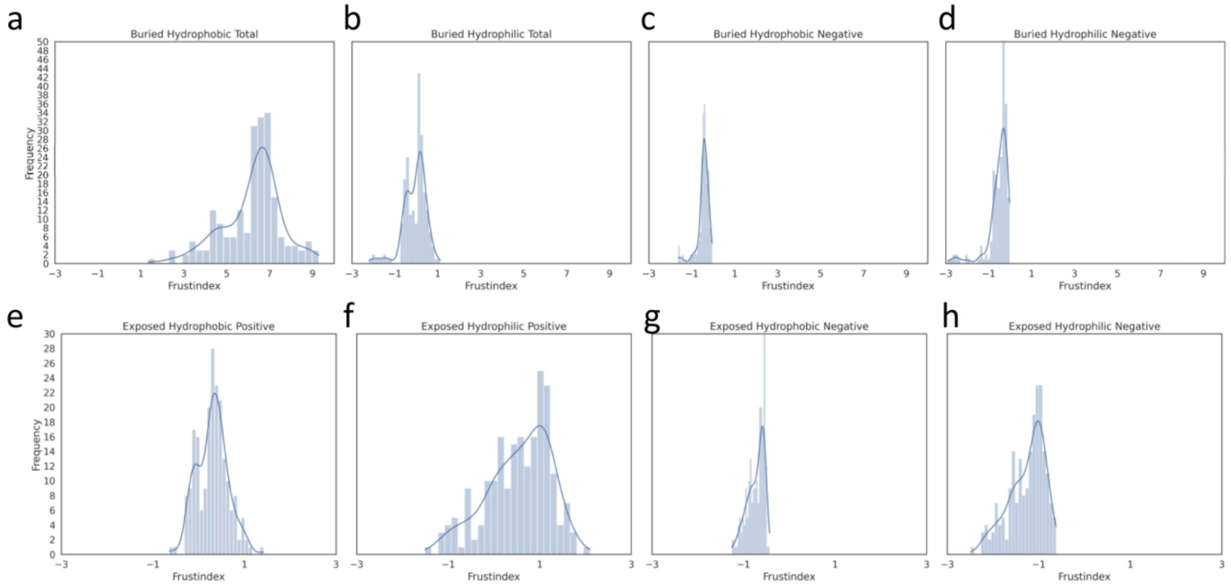


**Supplementary Fig. 1 | Configurational-frustration distributions across residue environments in small-molecule binding sites.** The analysis included 387 high-affinity-index protein–small-molecule complexes. Receptor residues were separated according to burial state (buried or exposed) and physicochemical class (hydrophobic or hydrophilic). Positive and negative configurational-frustration indices were summarized separately using their summed and mean values. Panels a–h show the corresponding distributions for the residue environments and frustration summaries indicated in each panel.

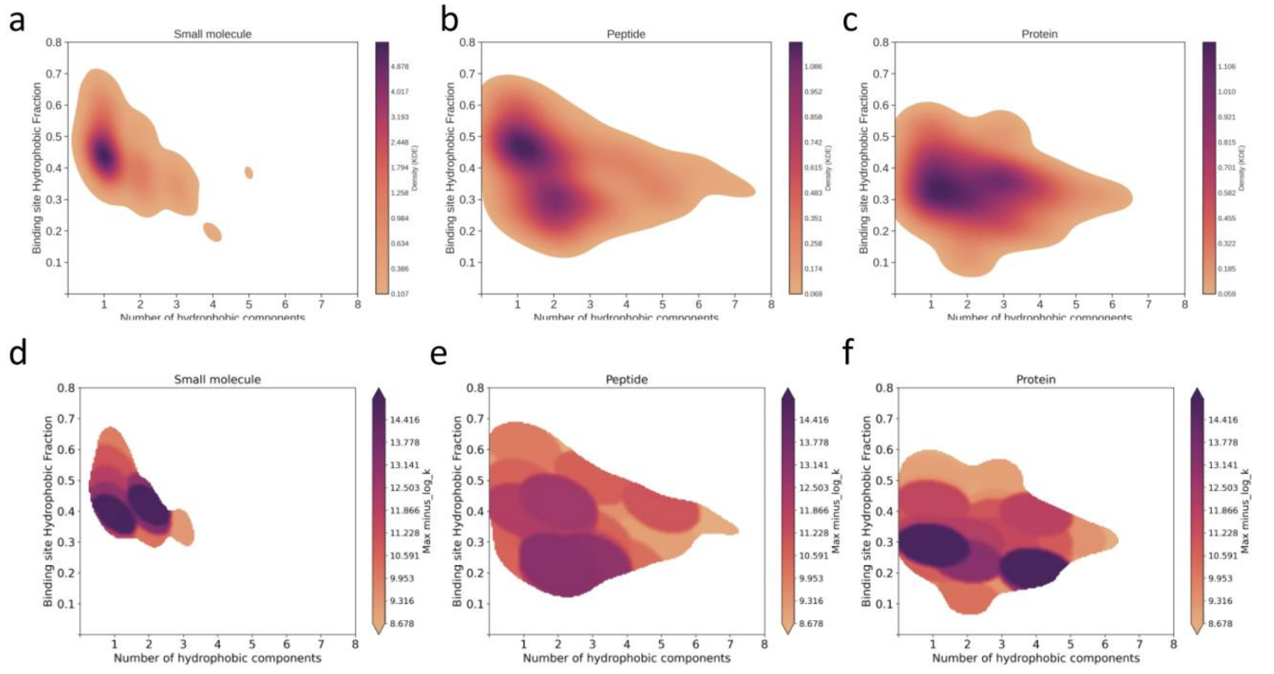


**Supplementary Fig. 2 | Joint organization of hydrophobic components in the high-affinity-index reference libraries.** Joint distributions of the number of connected hydrophobic components and the fraction of hydrophobic residues within the binding site are shown for protein–small-molecule (n = 387), protein–peptide (n = 186) and protein–protein (n = 246) complexes. Panels a–c correspond to small-molecule, peptide and protein binding sites, respectively, with colour indicating the estimated distribution density. Panels d–f show the corresponding small-molecule, peptide and protein distributions, respectively, coloured according to the maximum affinity index associated with each region of the distribution. The affinity index was defined as −log10(X [M]), where X denotes the reported Kd, Ki or IC50.

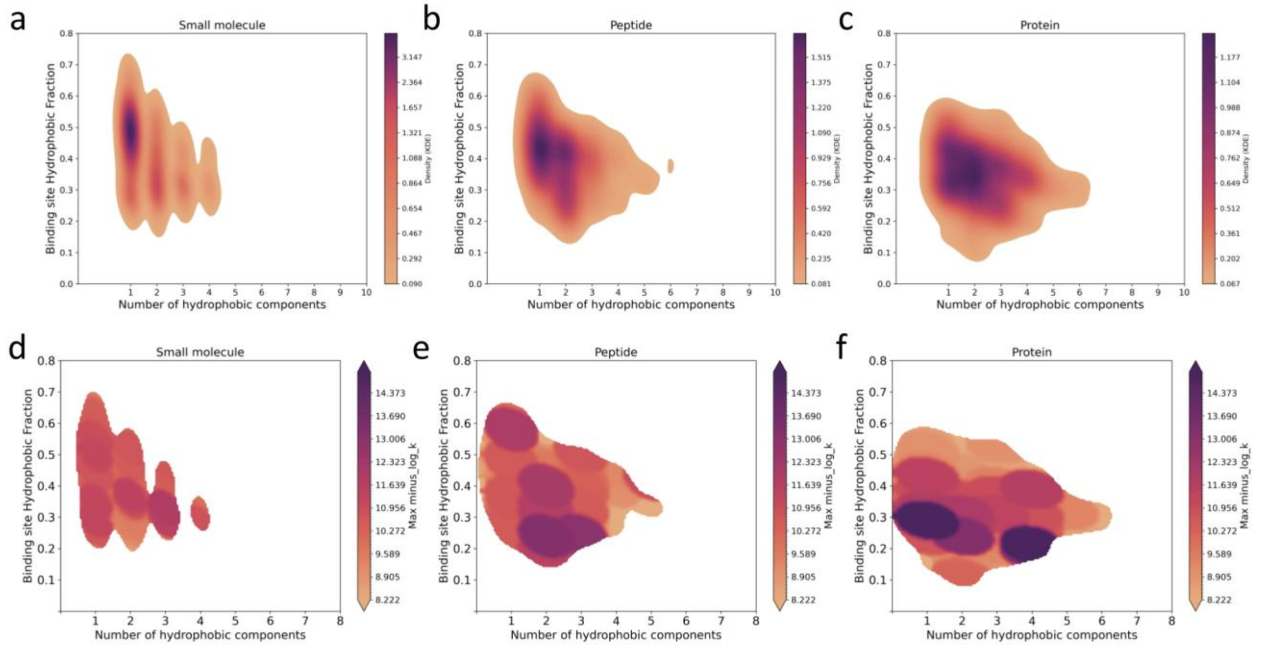


**Supplementary Fig. 3 | Joint organization of hydrophobic components in the affinity-unrestricted control libraries.** Joint distributions of the number of connected hydrophobic components and the fraction of hydrophobic residues within the binding site are shown for the affinity-unrestricted protein–small-molecule (n = 3,055), protein–peptide (n = 349) and protein–protein (n = 483) control libraries. Panels a–c correspond to small-molecule, peptide and protein binding sites, respectively, with colour indicating the estimated distribution density. Panels d–f show the corresponding small-molecule, peptide and protein distributions, respectively, coloured according to the maximum available affinity index associated with each region of the distribution. Affinity measurements were retained as metadata when available but were not used to select the control complexes.

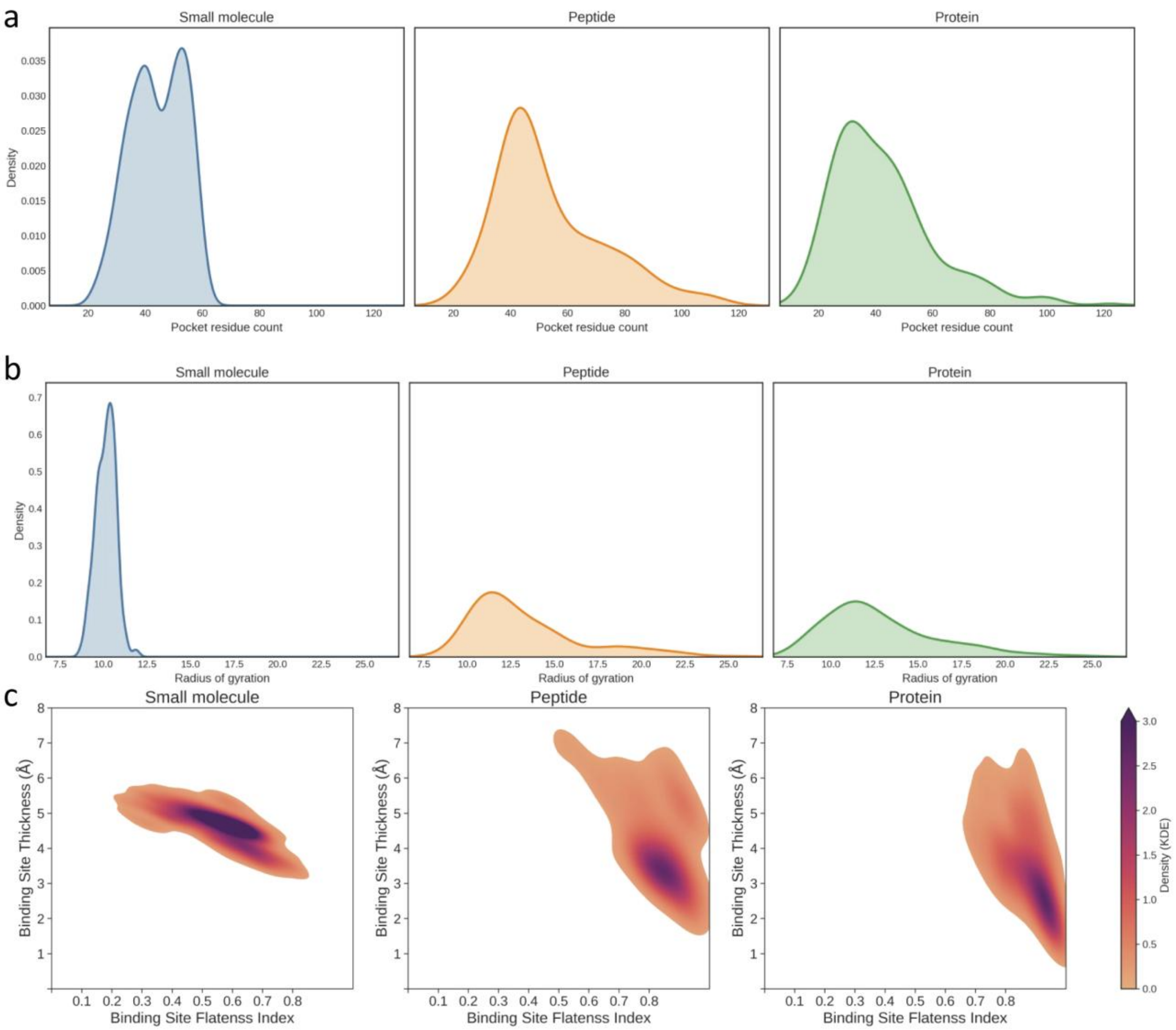


**Supplementary Fig. 4 | Binding-site size and shape distributions across molecular modalities.** a, Distributions of binding-site residue count for small-molecule, peptide and protein sites. b, Distributions of binding-site radius of gyration. c, Joint distributions of binding-site flatness index and thickness, coloured by density, for the three modalities.

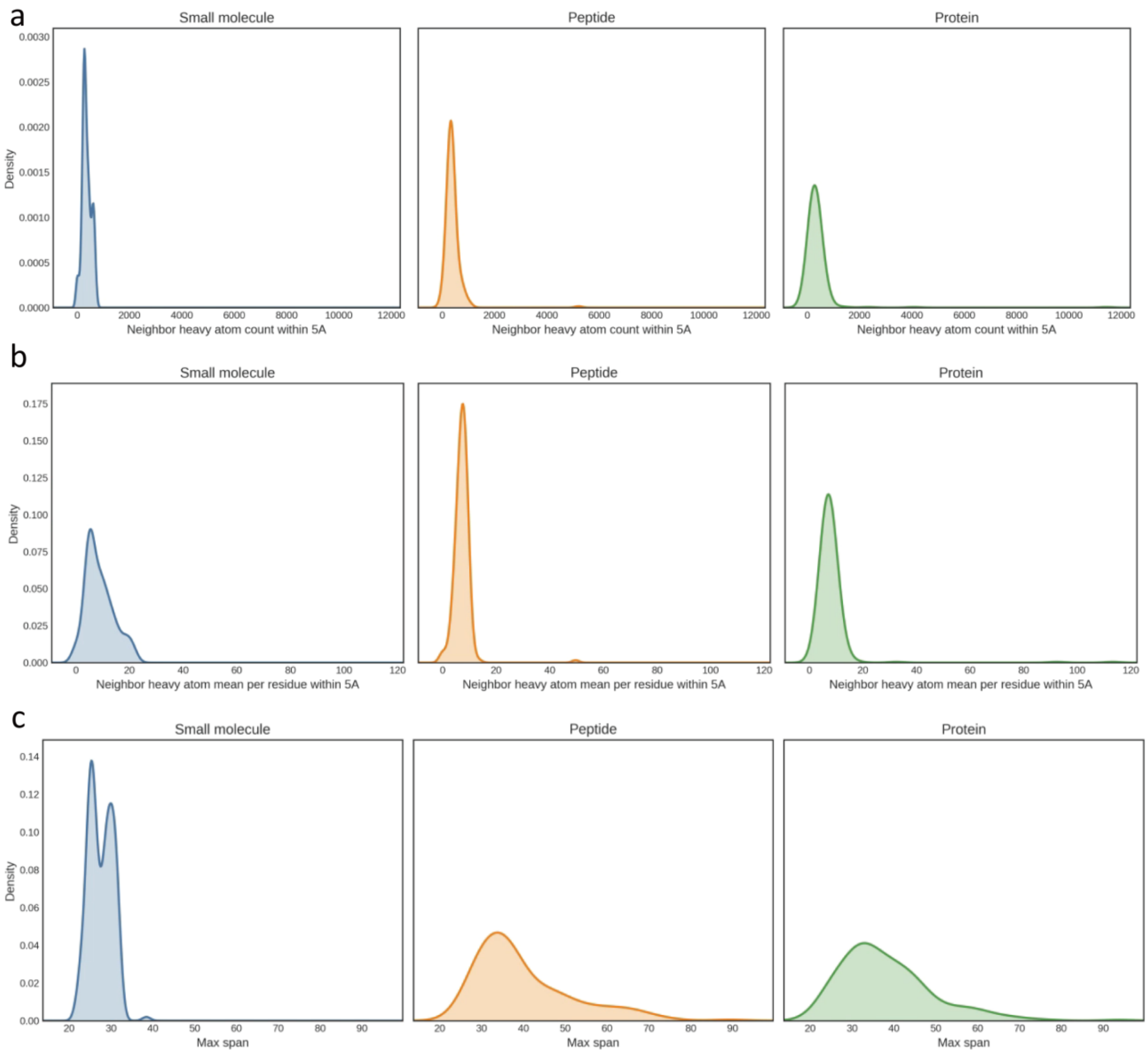


**Supplementary Fig. 5 | Local packing and spatial-span descriptors of binding sites.** a, Distributions of neighbouring heavy-atom counts within 5 Å for small-molecule, peptide-specific and protein binding sites. b, Mean neighbouring heavy-atom count per binding-site residue. All neighbouring-atom calculations included heavy atoms only. c, Maximum binding-site span, defined as the largest Euclidean distance between any pair of binding-site residue Cα atoms, for small-molecule, peptide-specific and protein binding sites.

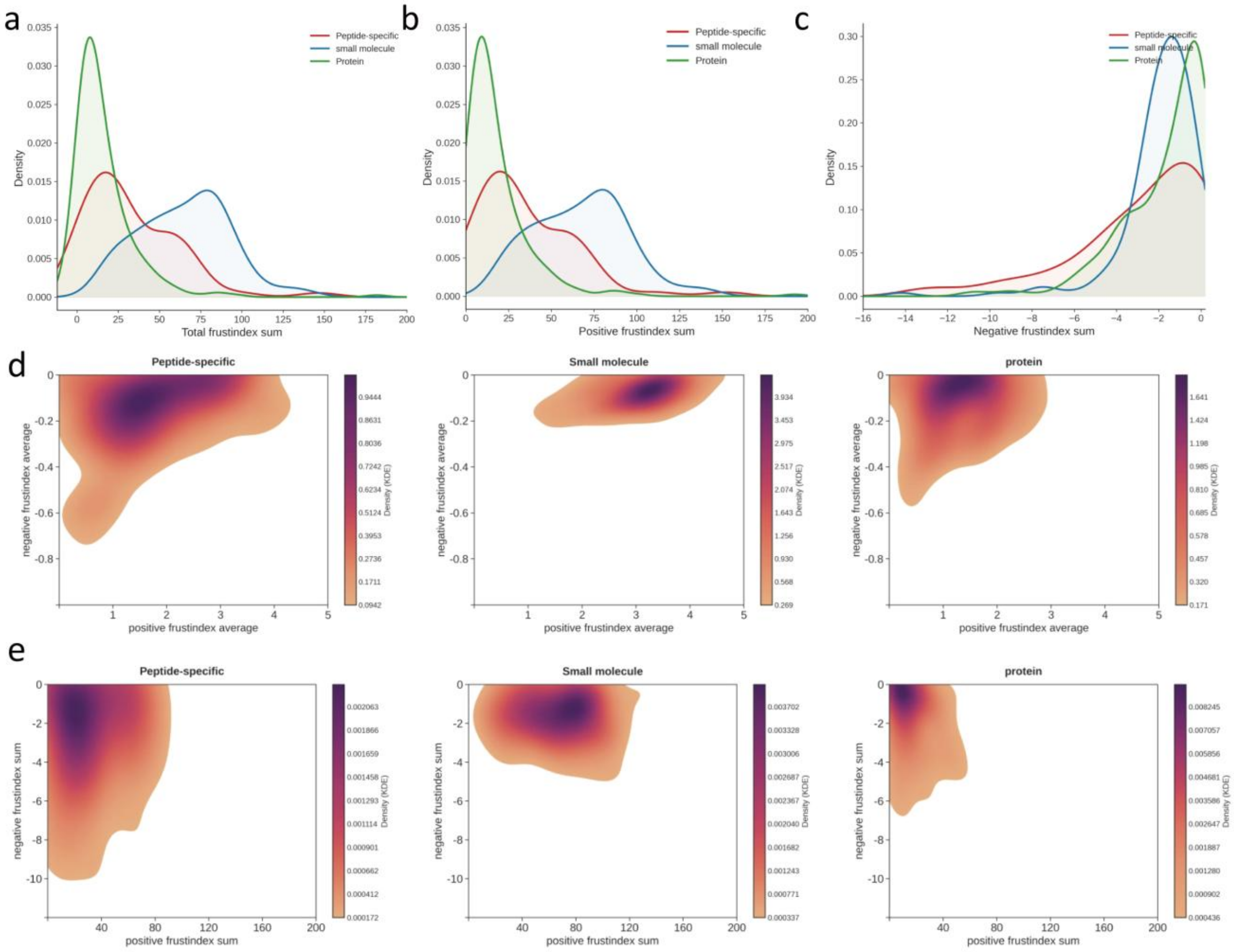


**Supplementary Fig. 6 | Configurational-frustration distributions across peptide-specific, small-molecule and protein binding sites.** All panels compare peptide-specific, small-molecule and protein binding sites. a–c, Density distributions of the summed total, positive and negative configurational-frustration indices, respectively. d, Joint distributions of the mean positive and negative configurational-frustration indices. e, Joint distributions of the summed positive and negative configurational-frustration indices.

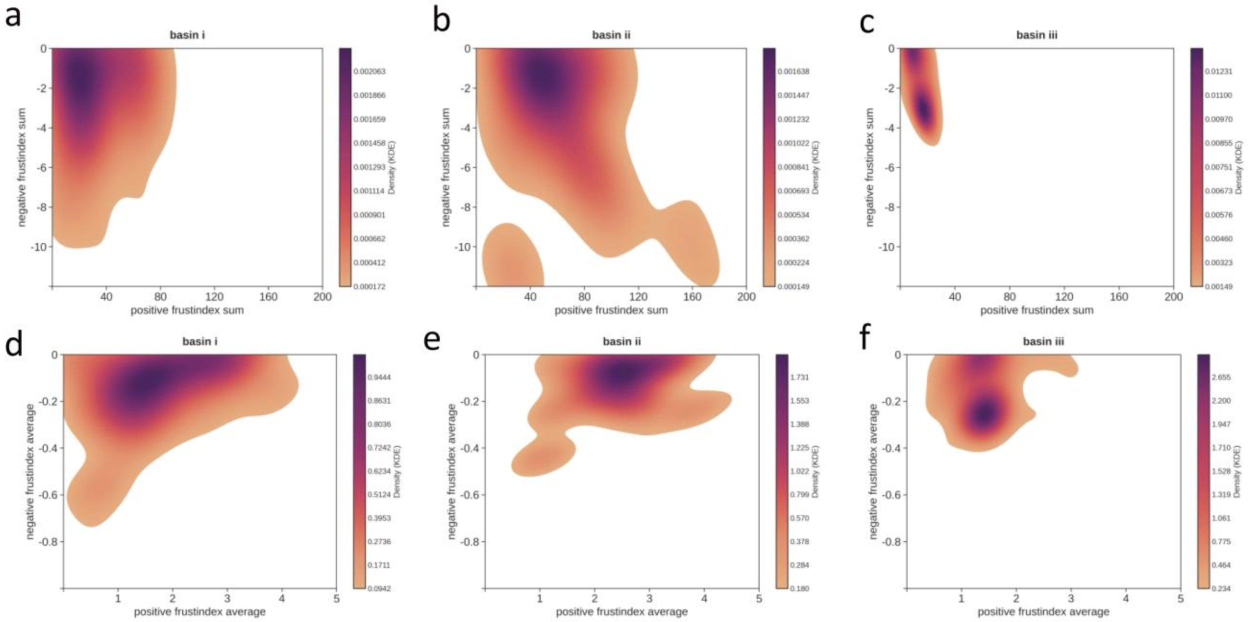


**Supplementary Fig. 7 | Configurational-frustration landscapes of the three peptide-recognition regimes.** a–c, Joint distributions of the summed positive and negative configurational-frustration indices for basin i (peptide-specific, n = 62), basin ii (small-molecule-like, n = 61) and basin iii (protein-like, n = 44), respectively. d–f, Corresponding joint distributions of the mean positive and negative configurational-frustration indices for basins i–iii, respectively. Recognition-regime assignments are defined in Fig. 2a.

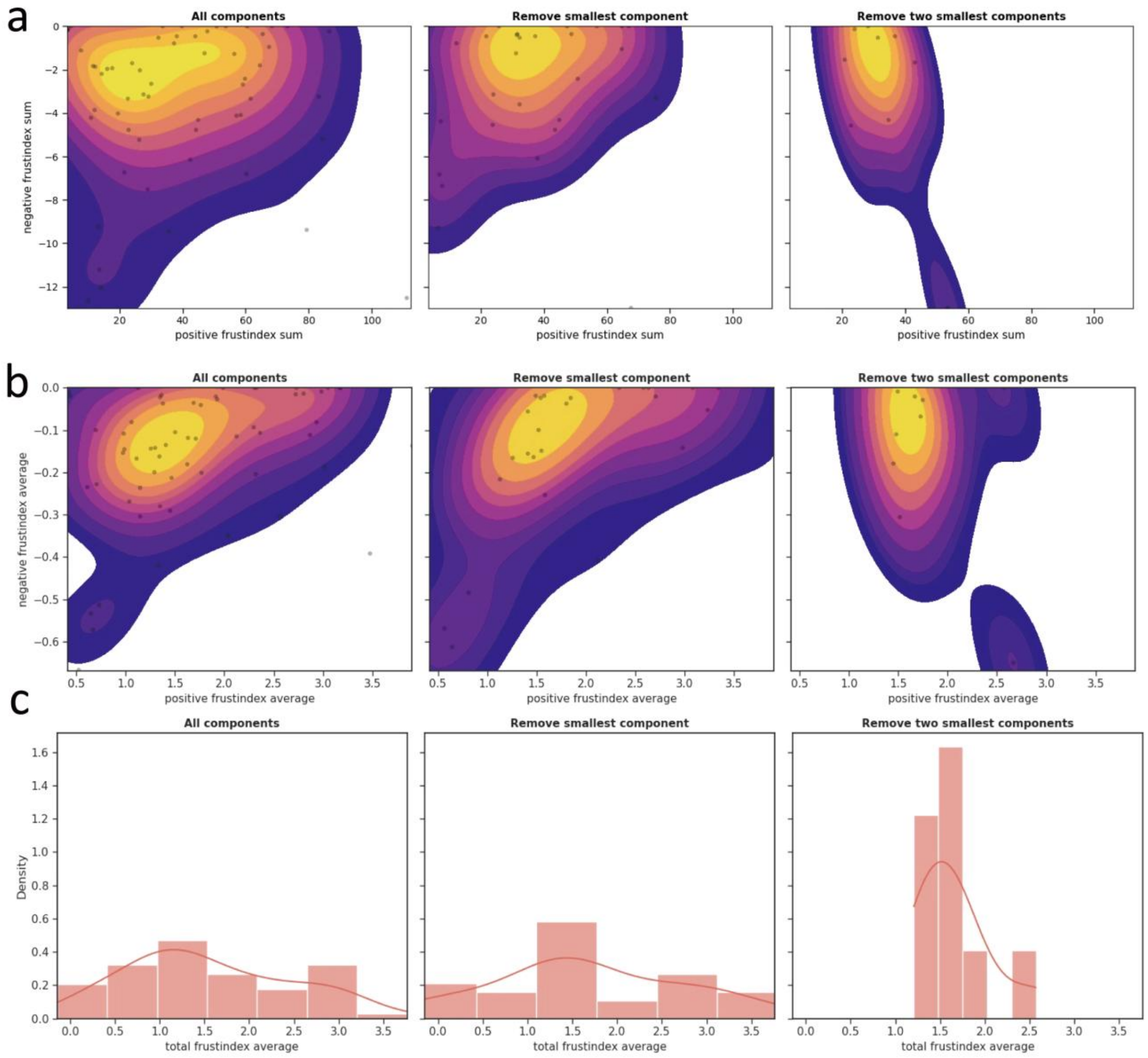


**Supplementary Fig. 8 | Effect of sequential component removal on frustration organization.** a, Joint distributions of positive and negative frustration sums with all components retained, after removal of the smallest component and after removal of the two smallest components. b, Corresponding distributions of average positive and negative frustration. c, Distributions of total average frustration under the three component-retention conditions.

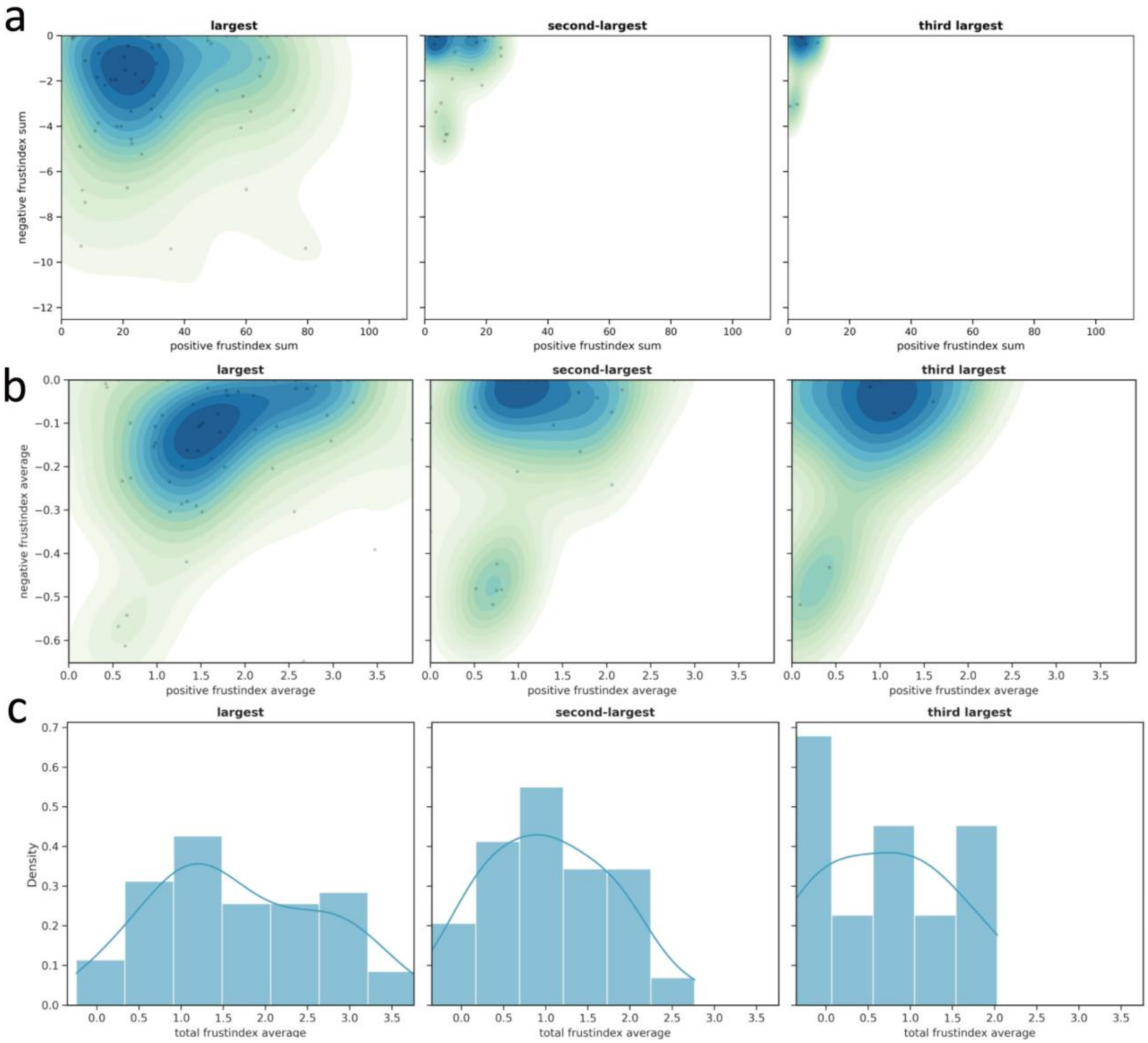


**Supplementary Fig. 9 | Frustration hierarchy among the three largest hydrophobic components.** a, Positive and negative frustration-sum landscapes of the largest, second-largest and third-largest components. b, Corresponding landscapes of average positive and negative frustration. c, Distributions of total average frustration for the three component ranks.

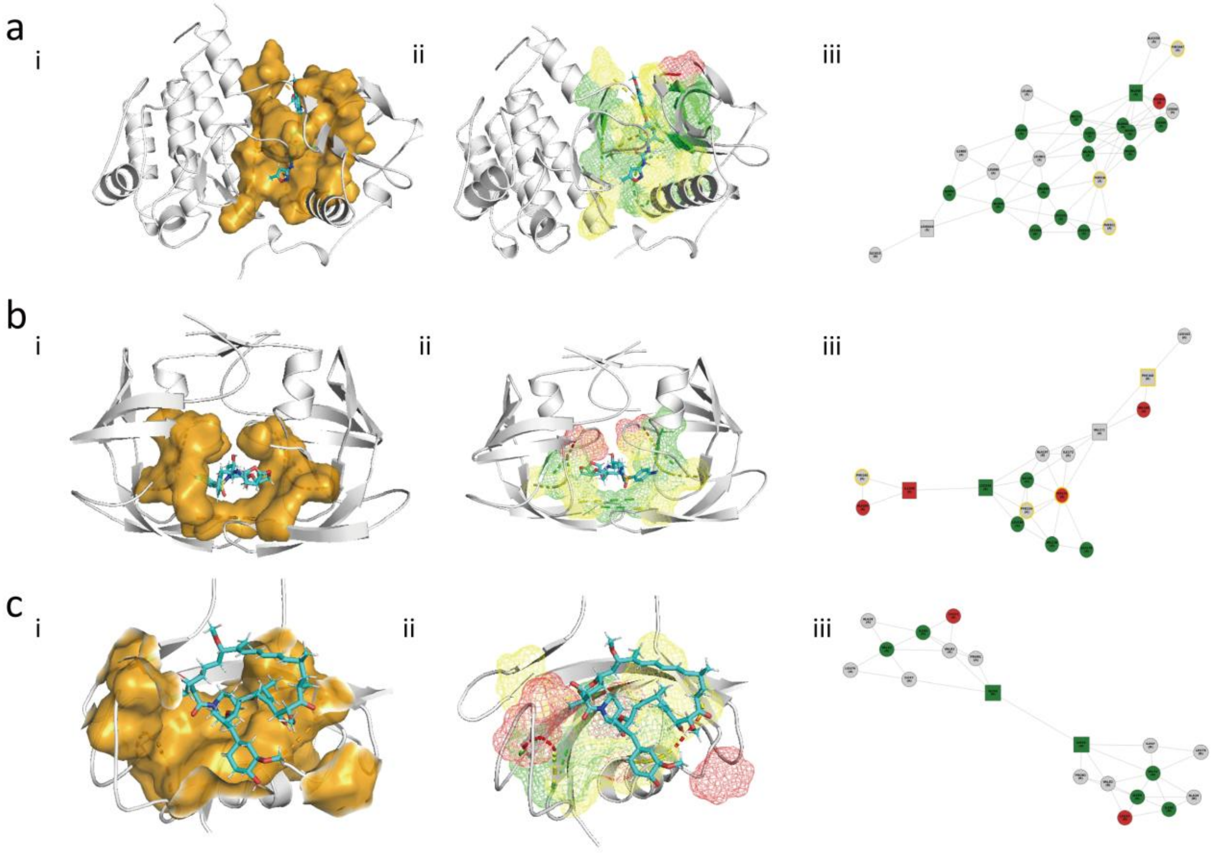


**Supplementary Fig. 10 | Representative small-molecule binding-site architectures.** Rows a–c show representative protein–small-molecule complexes with PDB IDs 4ASE, 1PBK and 6OOT, respectively. In each row, i shows the receptor as a white cartoon, the small-molecule ligand as cyan sticks and the connected hydrophobic components as orange surfaces; ii maps the spatial distribution of configurational frustration within the largest hydrophobic component, with green, yellow and red indicating minimally frustrated, intermediate and highly frustrated regions, respectively; and iii shows the corresponding hydrophobic-residue connectivity graph.

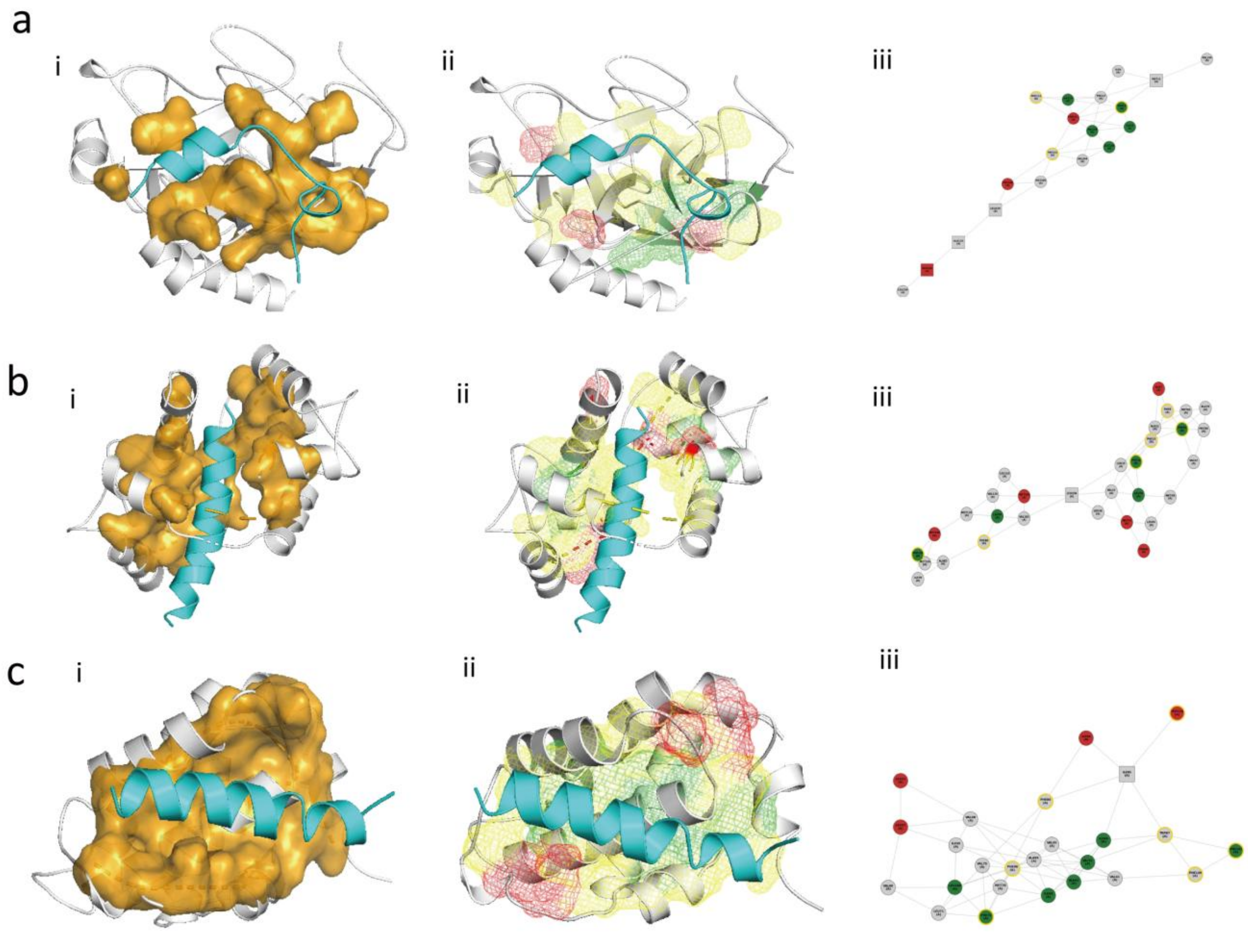


**Supplementary Fig. 11 | Representative basin-ii peptide-binding architectures.** Rows a–c show representative small-molecule-like peptide-binding complexes from basin ii with PDB IDs 1A3B, 3DVM and 5UUL, respectively. In each row, i shows the receptor as a white cartoon, the peptide as a cyan cartoon and the connected hydrophobic components as orange surfaces; ii maps the spatial distribution of configurational frustration within the largest hydrophobic component, with green, yellow and red indicating minimally frustrated, intermediate and highly frustrated regions, respectively; and iii shows the corresponding hydrophobic-residue connectivity graph.

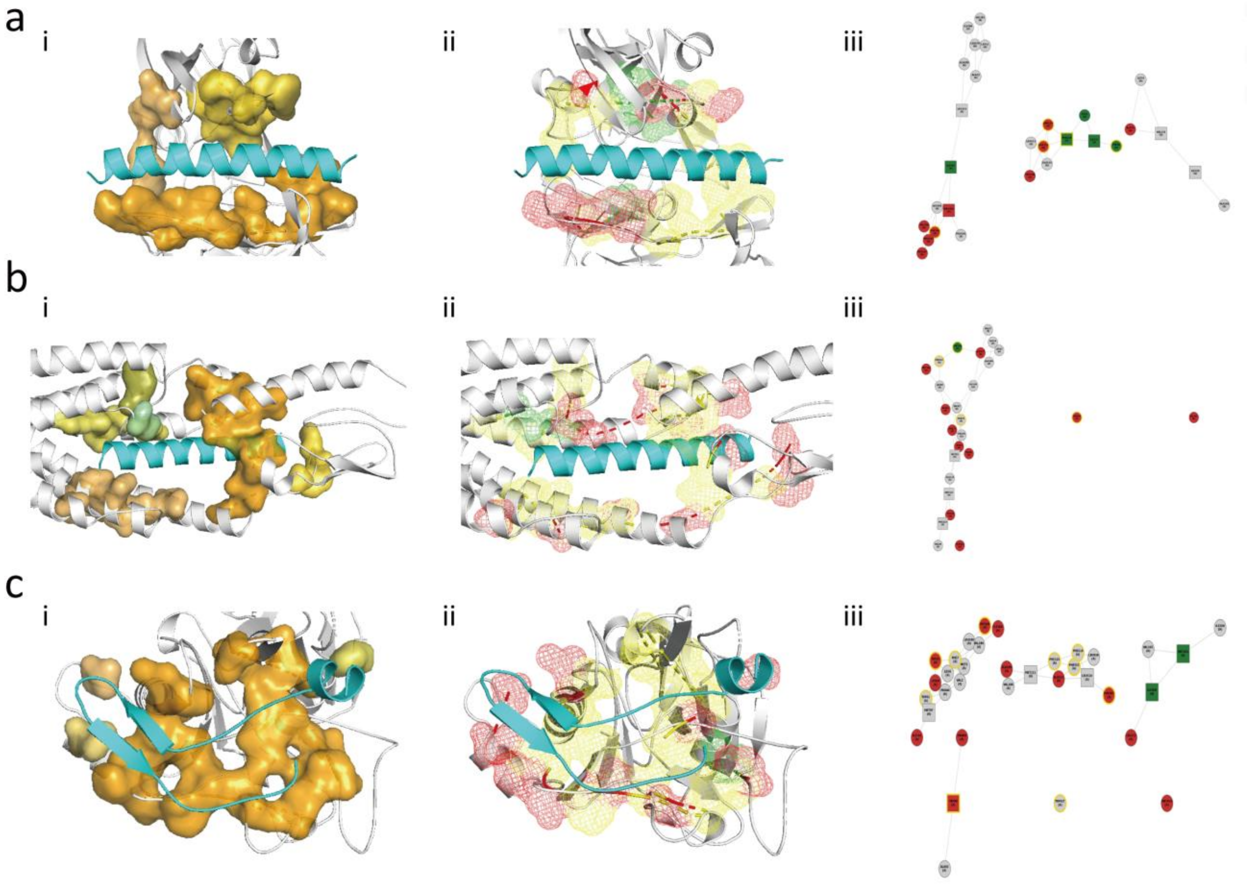


**Supplementary Fig. 12 | Representative basin-i peptide-specific binding architectures.** Rows a–c show representative peptide-specific complexes from basin i with PDB IDs 1DPJ, 5YQZ and 3ZWZ, respectively. In each row, i shows the receptor as a white cartoon, the peptide as a cyan cartoon and the connected hydrophobic components as orange surfaces; ii maps the spatial distribution of configurational frustration within the largest hydrophobic component, with green, yellow and red indicating minimally frustrated, intermediate and highly frustrated regions, respectively; and iii shows the corresponding hydrophobic-residue connectivity graph.

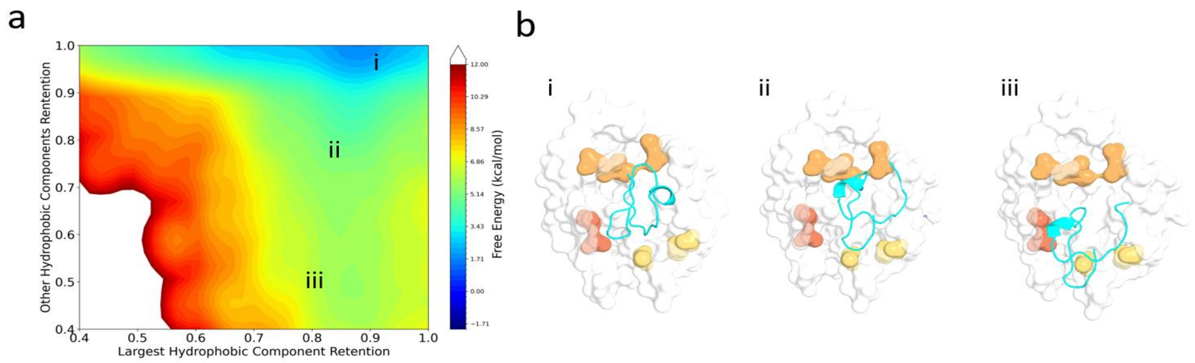


**Supplementary Fig. 13 | Free-energy landscape of hydrophobic-component retention.** a, Two-dimensional free-energy surface for the representative peptide–receptor complex 1MCV, projected onto retention of the largest hydrophobic component and the remaining components. b, Representative conformations corresponding to states i–iii marked on the landscape. The largest (C1), second-largest (C2) and third-largest (C3) hydrophobic components are coloured red, orange and yellow, respectively. State i retains engagement with C1, C2 and C3; state ii retains engagement with C1 and C2; and state iii retains engagement with C2 and C3.

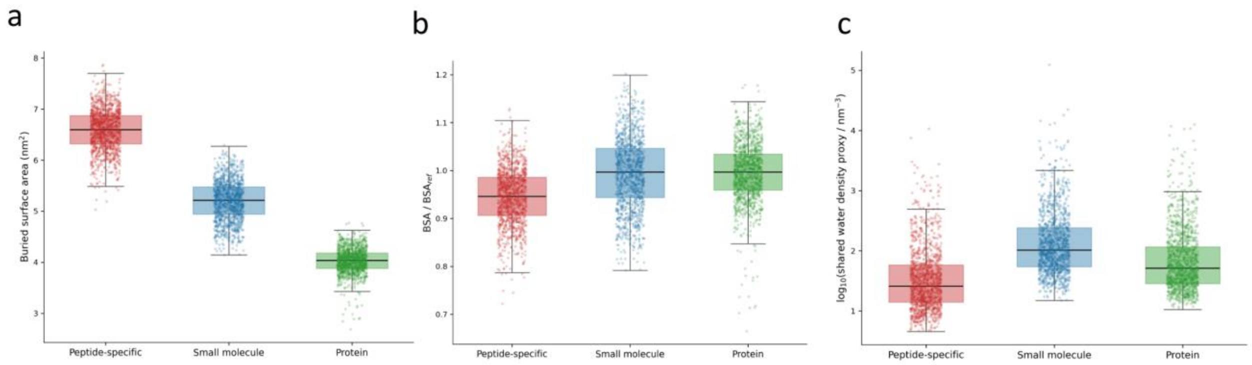


**Supplementary Fig. 14 | Interfacial geometry and hydration across molecular modalities.** Distributions are shown for the peptide-specific complex 1DPJ, the protein–small-molecule complex 1MUI and the protein–protein complex 1A22. a, Buried surface area (BSA), reported in $nm^2$. b, BSA ratio, calculated by normalizing the value in each frame to that in the first analysed frame. c, Shared-water density proxy, calculated from water oxygen atoms simultaneously located within 0.35 nm of pocket and binder heavy atoms and displayed on a log10 scale.

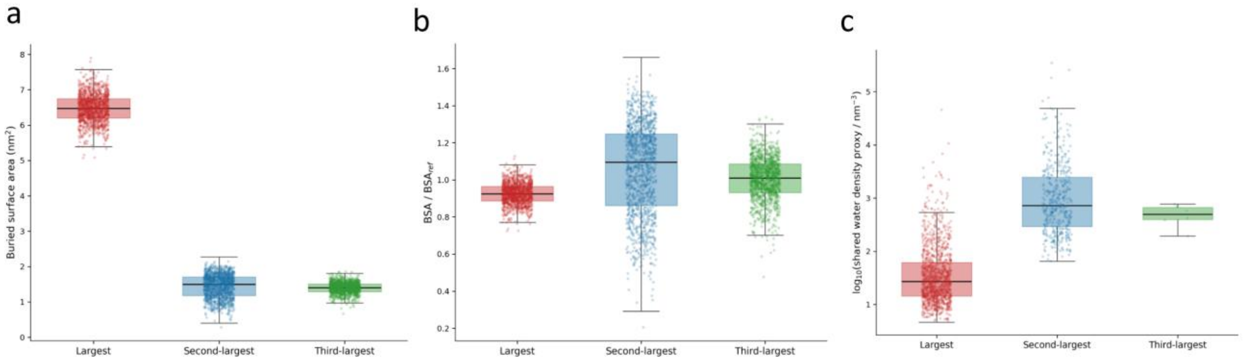


**Supplementary Fig. 15 | Component-resolved geometry and hydration within a peptide-specific binding site.** Distributions are shown for the largest (C1), second-largest (C2) and third-largest (C3) hydrophobic components of the peptide-specific complex 1DPJ. a, Buried surface area (BSA), reported in $nm^2$. b, BSA ratio, calculated by normalizing the value in each frame to that in the first analysed frame. c, Shared-water density proxy, calculated from water oxygen atoms simultaneously located within 0.35 nm of the component and peptide heavy atoms and displayed on a log10 scale.

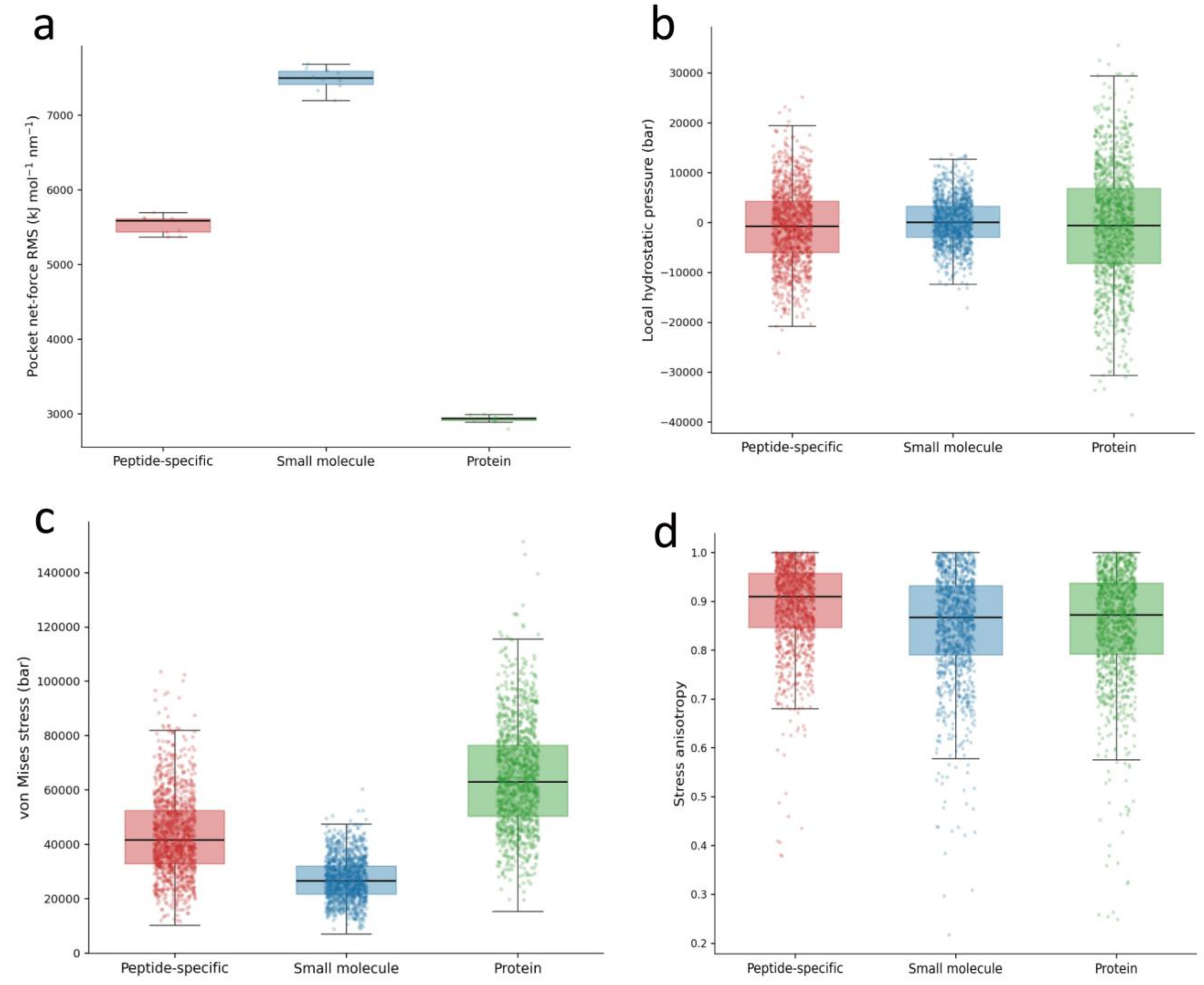


**Supplementary Fig. 16 | Mechanical descriptors across molecular modalities.** Distributions are shown for the peptide-specific complex 1DPJ, the protein–small-molecule complex 1MUI and the protein–protein complex 1A22. a, Pocket net-force root-mean-square magnitude, calculated over consecutive 500-frame windows and reported in kJ $mol^{-1}$ $nm^{-1}$. b, Local hydrostatic pressure, reported in bar. c, von Mises stress, reported in bar. d, Dimensionless stress anisotropy.

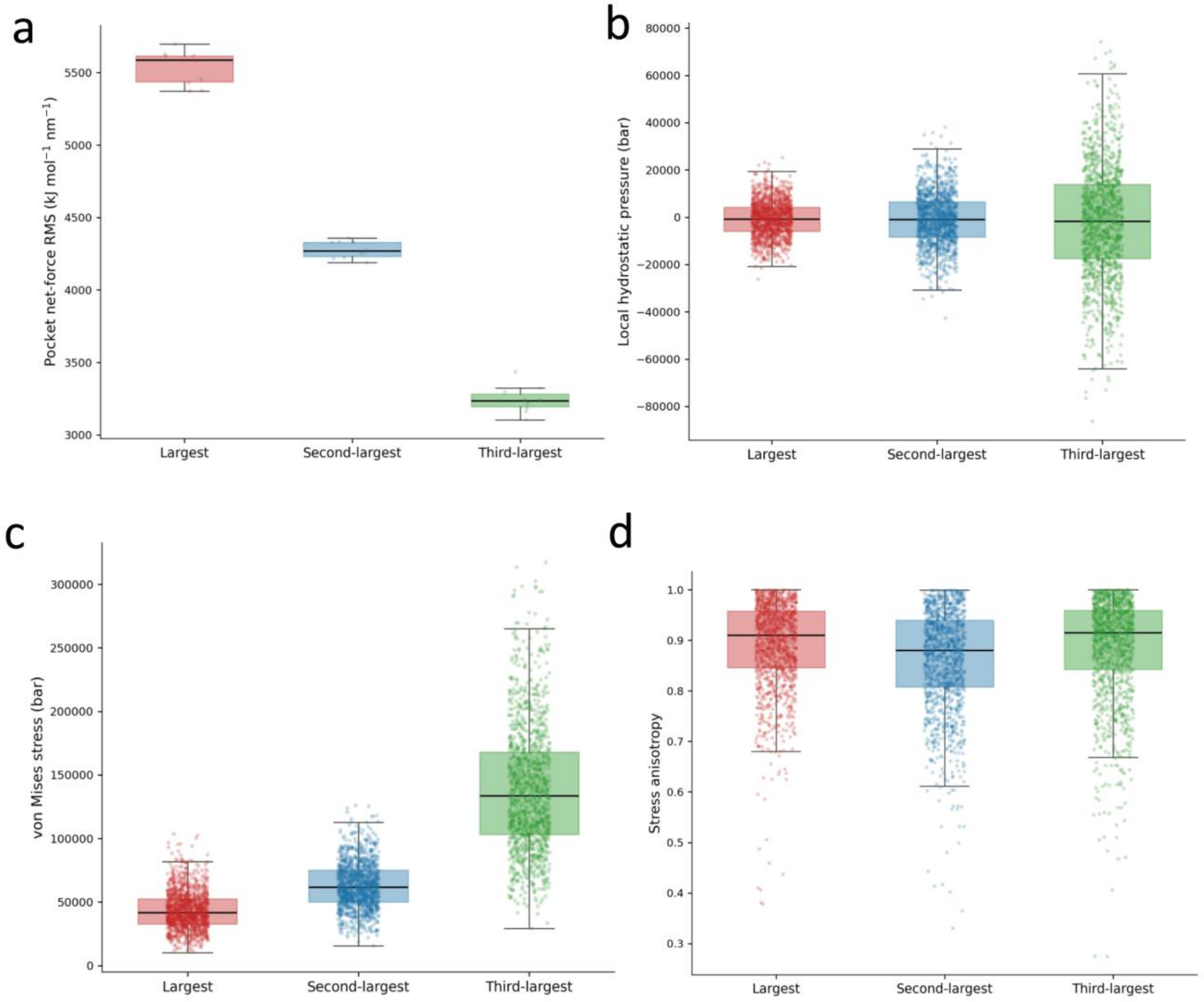


**Supplementary Fig. 17 | Component-resolved mechanical descriptors within a peptide-specific binding site.** Distributions are shown for the largest (C1), second-largest (C2) and third-largest (C3) hydrophobic components of the peptide-specific complex 1DPJ. a, Component net-force root-mean-square magnitude, calculated over consecutive 500-frame windows and reported in kJ mol$^{-1}$ nm$^{-1}$. b, Local hydrostatic pressure, reported in bar. c, von Mises stress, reported in bar. d, Dimensionless stress anisotropy.

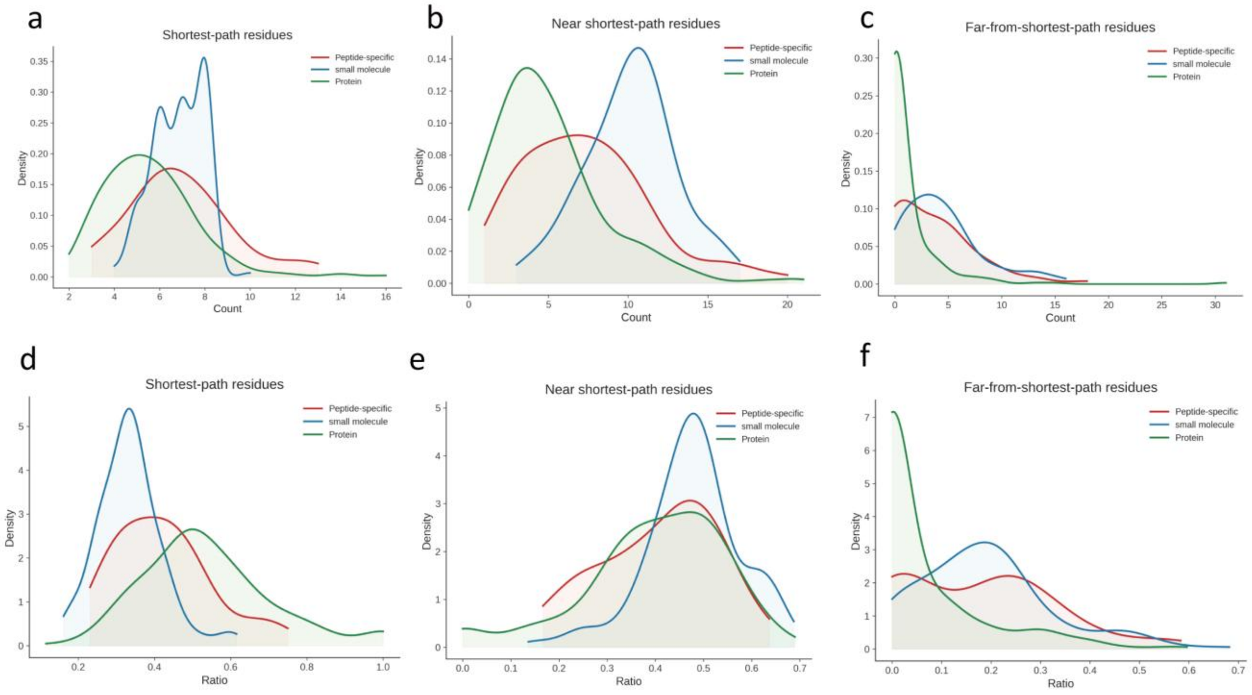


**Supplementary Fig. 18 | Topological organization of shortest-path and peripheral residues.** a–c, Distributions of residue counts assigned to the shortest-path, near-shortest-path and far-from-shortest-path regions for peptide-specific, small-molecule and protein binding sites. d–f, Corresponding distributions expressed as fractions of binding-site residues.

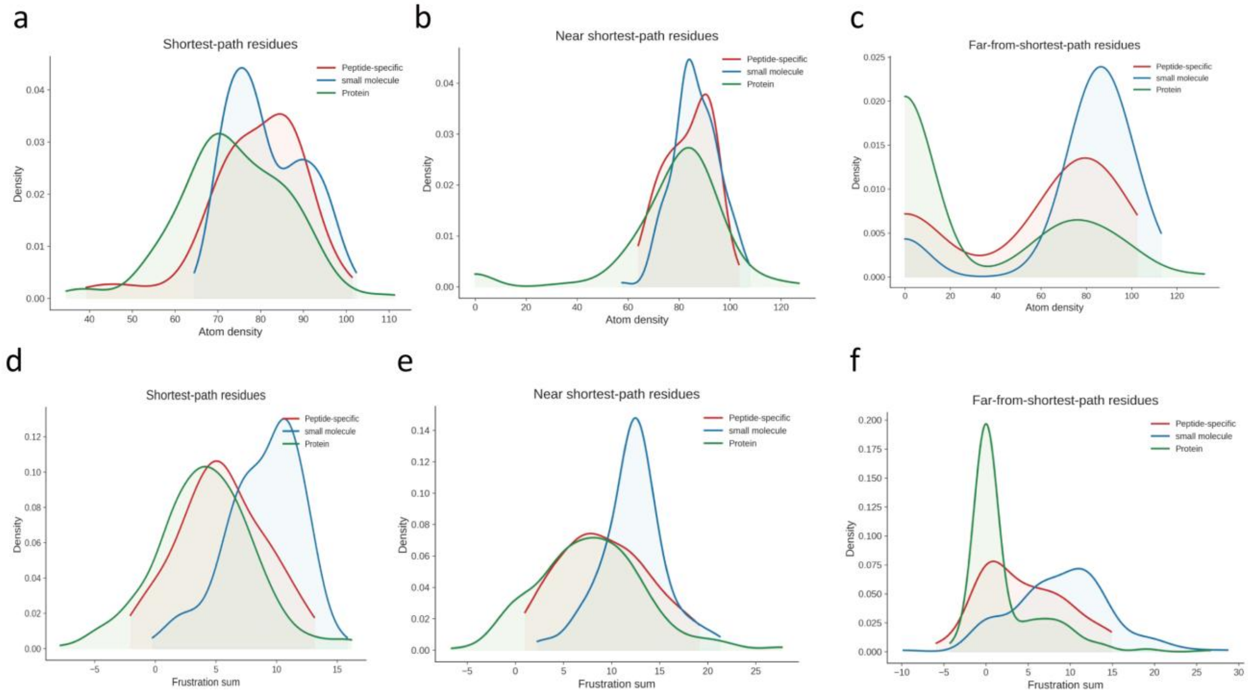


**Supplementary Fig. 19 | Packing and energetic properties across topological residue classes.** a–c, Atom-density distributions for shortest-path, near-shortest-path and far-from-shortest-path residues. d–f, Corresponding distributions of residue-level frustration sums for peptide-specific, small-molecule and protein sites.

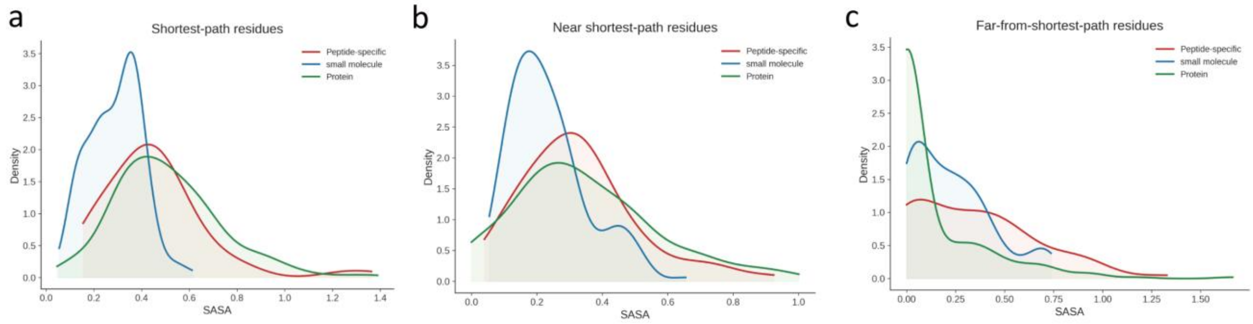


**Supplementary Fig. 20 | Solvent accessibility across topological residue classes.** Distributions of solvent-accessible surface area for a, shortest-path residues; b, near-shortest-path residues; and c, far-from-shortest-path residues in peptide-specific, small-molecule and protein binding sites.

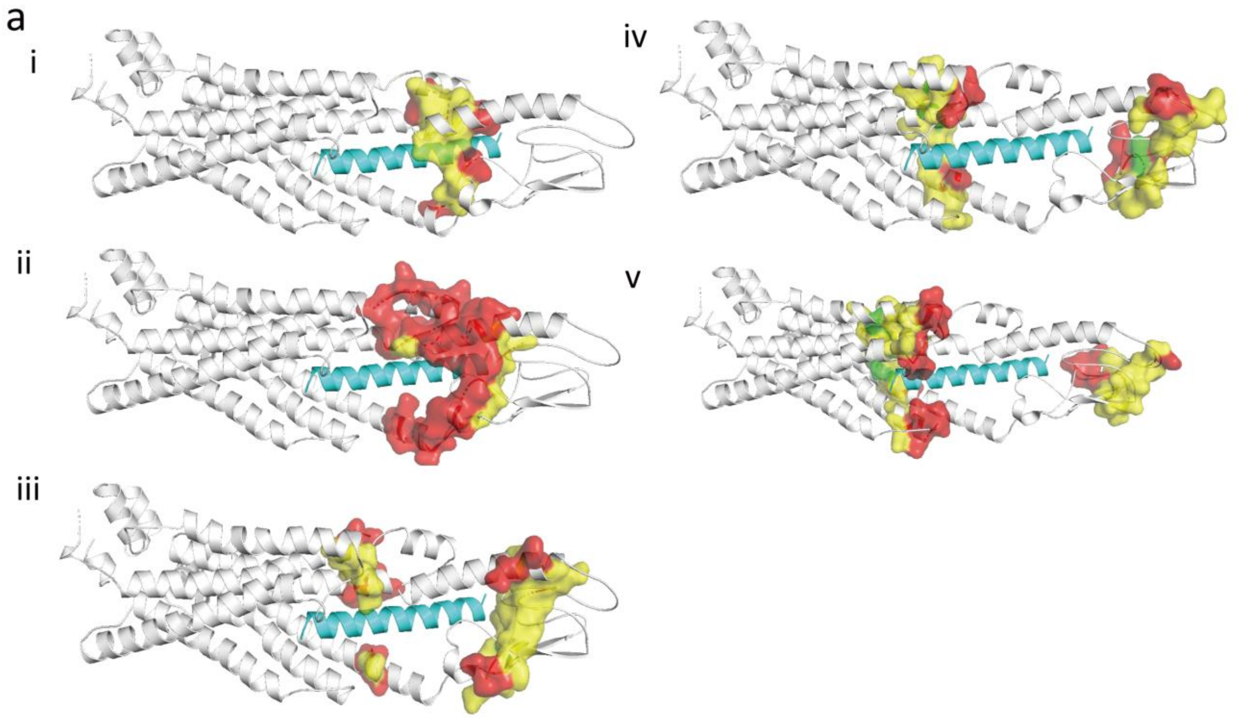


**Supplementary Fig. 21 | Representative receptor-shell organization in a peptide-specific complex.** Representative views of the peptide-specific receptor–peptide complex 5YQZ showing residues assigned to successive receptor shells. Panels i–v show S0, S1, S2, S3 and S5, respectively. The receptor is shown in white and the peptide in cyan, with the indicated shell residues highlighted according to the colours used in the structures.

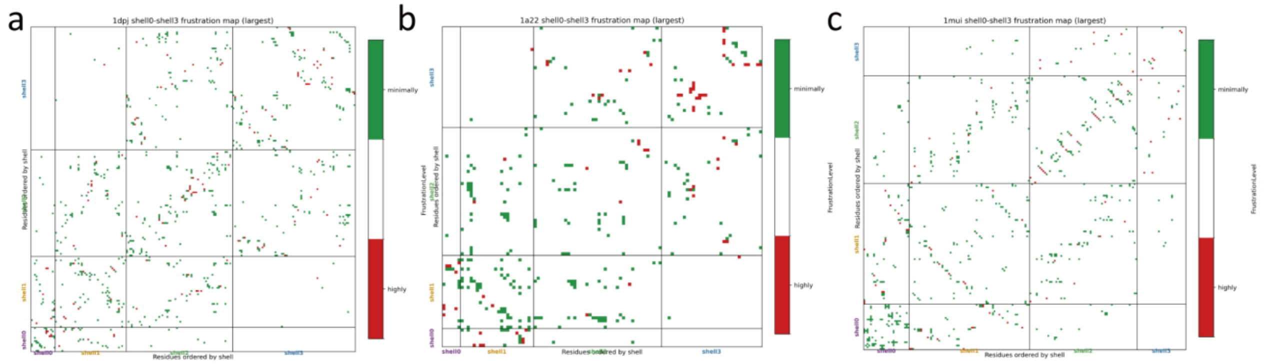


**Supplementary Fig. 22 | Shell-resolved maps of configurational-frustration changes.** a–c, Configurational-frustration change maps for the representative peptide-specific complex 1DPJ, protein–small-molecule complex 1MUI and protein–protein complex 1A22, respectively. Each map shows the organization of receptor shells S0, S1 and S2. Green and red marks denote changes in the directions indicated by the corresponding colour keys.

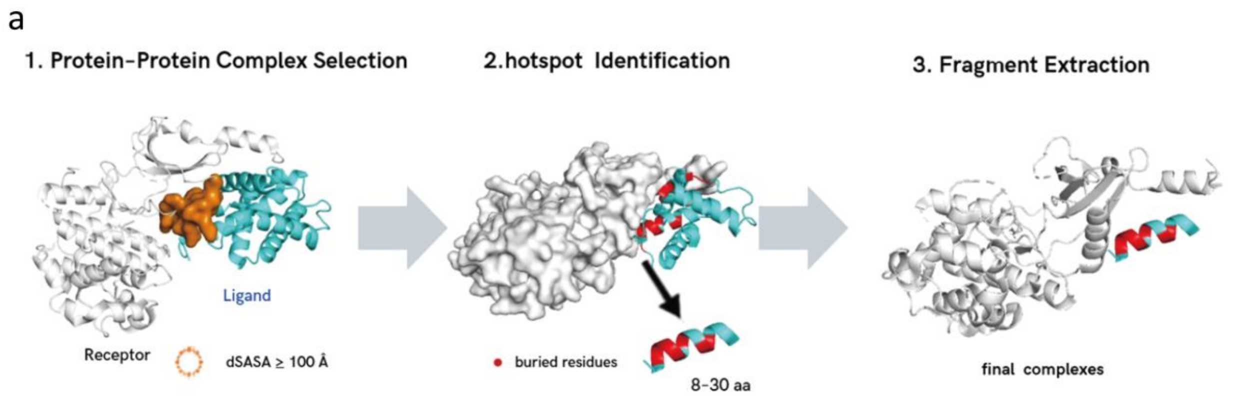


**Supplementary Fig. 23 | Geometry-based extraction of peptide-like fragments from protein interfaces.** Schematic of the conventional augmentation workflow comprising protein–protein complex selection, hotspot identification from buried interface residues and extraction of an 8–30-residue interacting fragment to form a receptor–peptide complex.

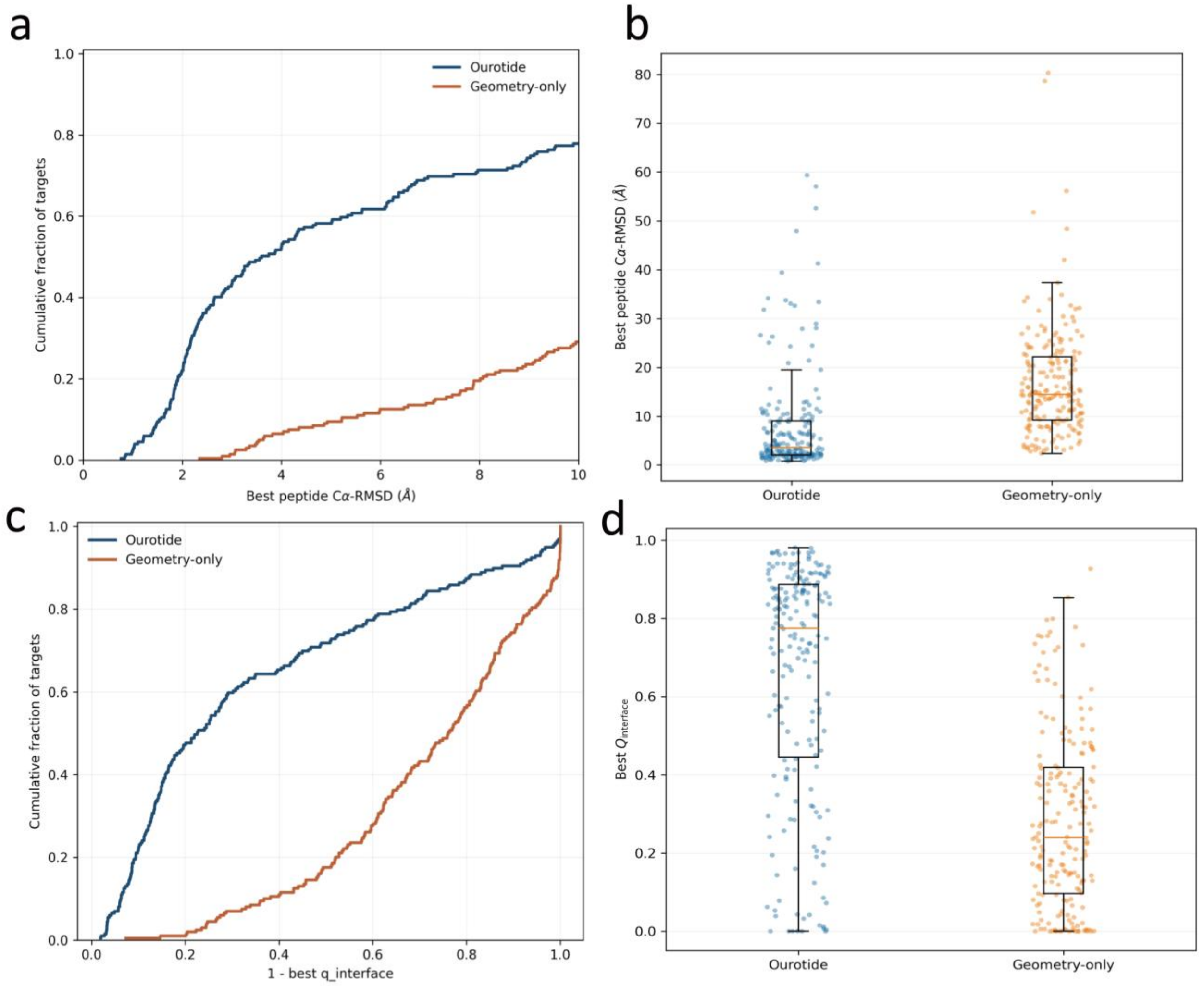


**Supplementary Fig. 24 | Effect of physics-guided augmentation on structure recovery.** Comparison of Ourotide trained with the physics-guided dataset and the geometry-only augmentation control. a, Cumulative distribution of best peptide Cα-RMSD. b, Distribution of best peptide Cα-RMSD. c, Cumulative distribution of interface-recovery error expressed as 1 − best Qinterface. d, Distribution of best Qinterface.

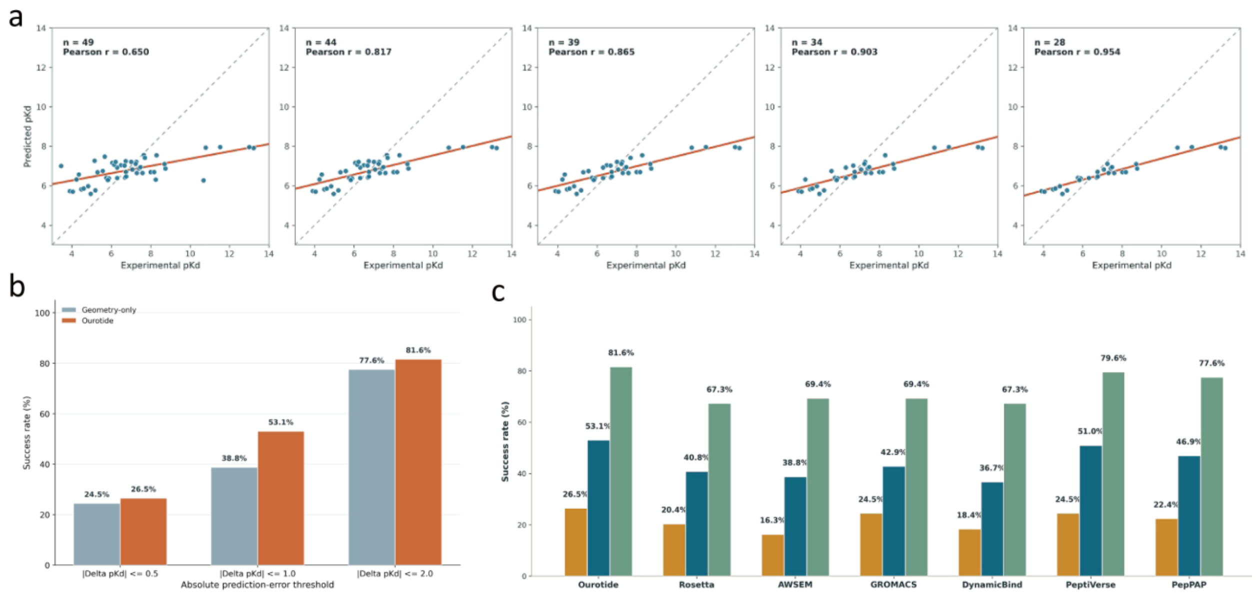


**Supplementary Fig. 25 | Effect of physics-guided augmentation on affinity prediction.** a, Predicted versus experimental pKd across the five evaluation folds, with the number of available complexes and Pearson correlation shown in each panel. b, Success rates of geometry-only and physics-guided Ourotide models at the indicated absolute-error thresholds. c, Success rates of the seven evaluated affinity-prediction methods.

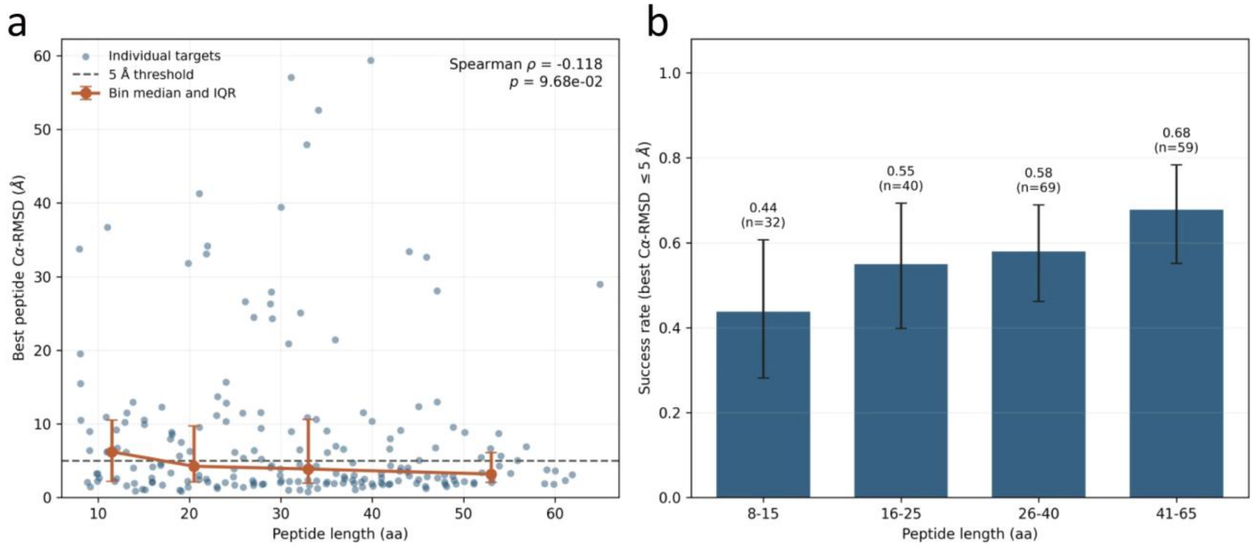


**Supplementary Fig. 26 | Dependence of structural recovery on peptide length.** a, Best peptide Cα-RMSD as a function of peptide length; points denote individual targets and binned summaries show the median and interquartile range. The dashed line marks the 5-Å threshold. b, Best-of-five Cα-RMSD success rate in the indicated peptide-length bins, with bin counts shown above the bars.

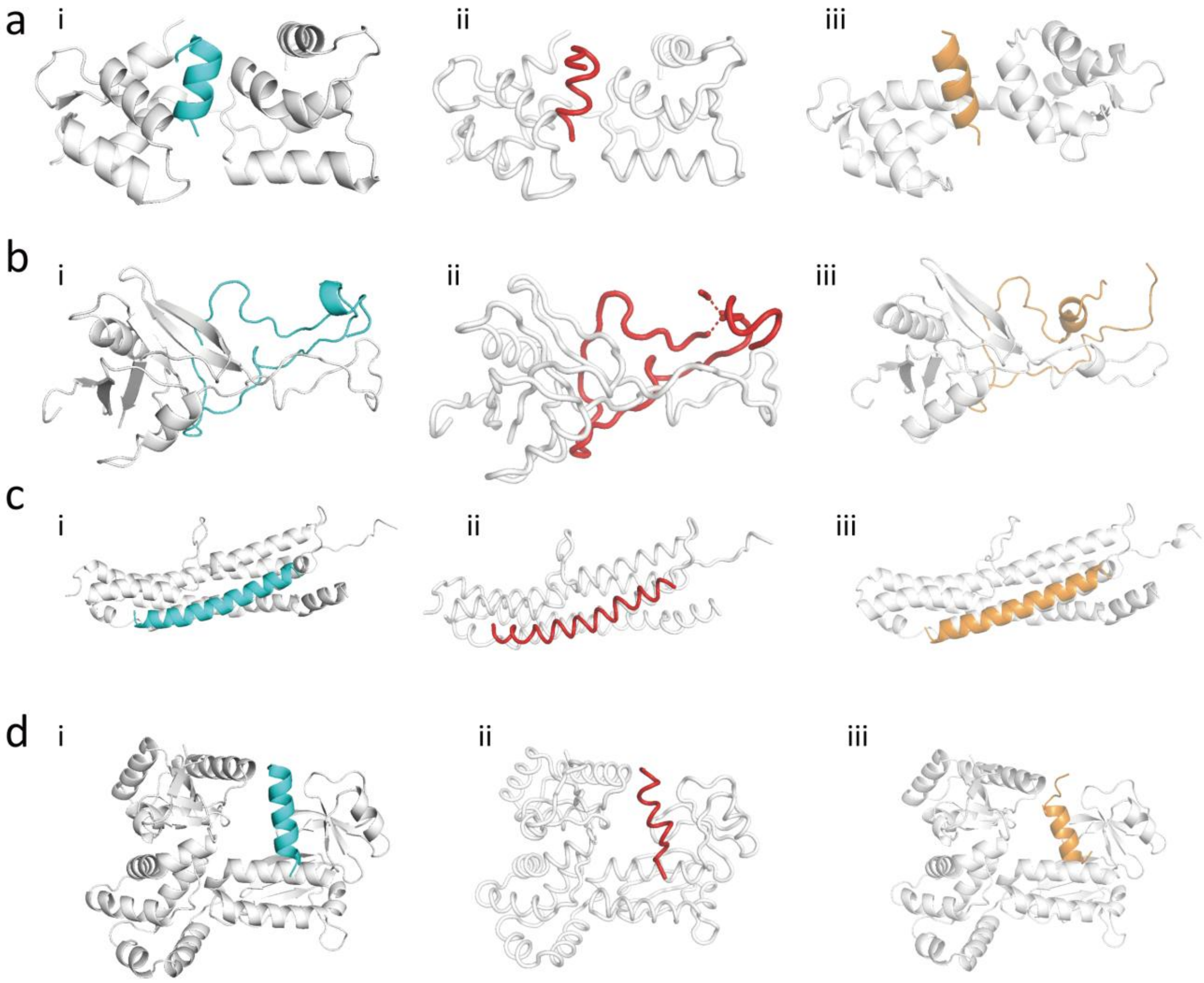


**Supplementary Fig. 27 | Additional examples of receptor-bound peptide structure prediction.** a–d, Representative receptor–peptide complexes with PDB IDs 4LZX, 5THP, 3JSE and 6CH2, respectively. For each complex, i shows the experimentally determined structure, ii shows the Ourotide prediction and iii shows the AlphaFold 3 prediction. Receptors are shown in white, whereas the experimental, Ourotide-predicted and AlphaFold 3-predicted peptides are shown in cyan, red and orange, respectively. The prediction with the lowest peptide Cα-RMSD among five candidates was selected for each method.

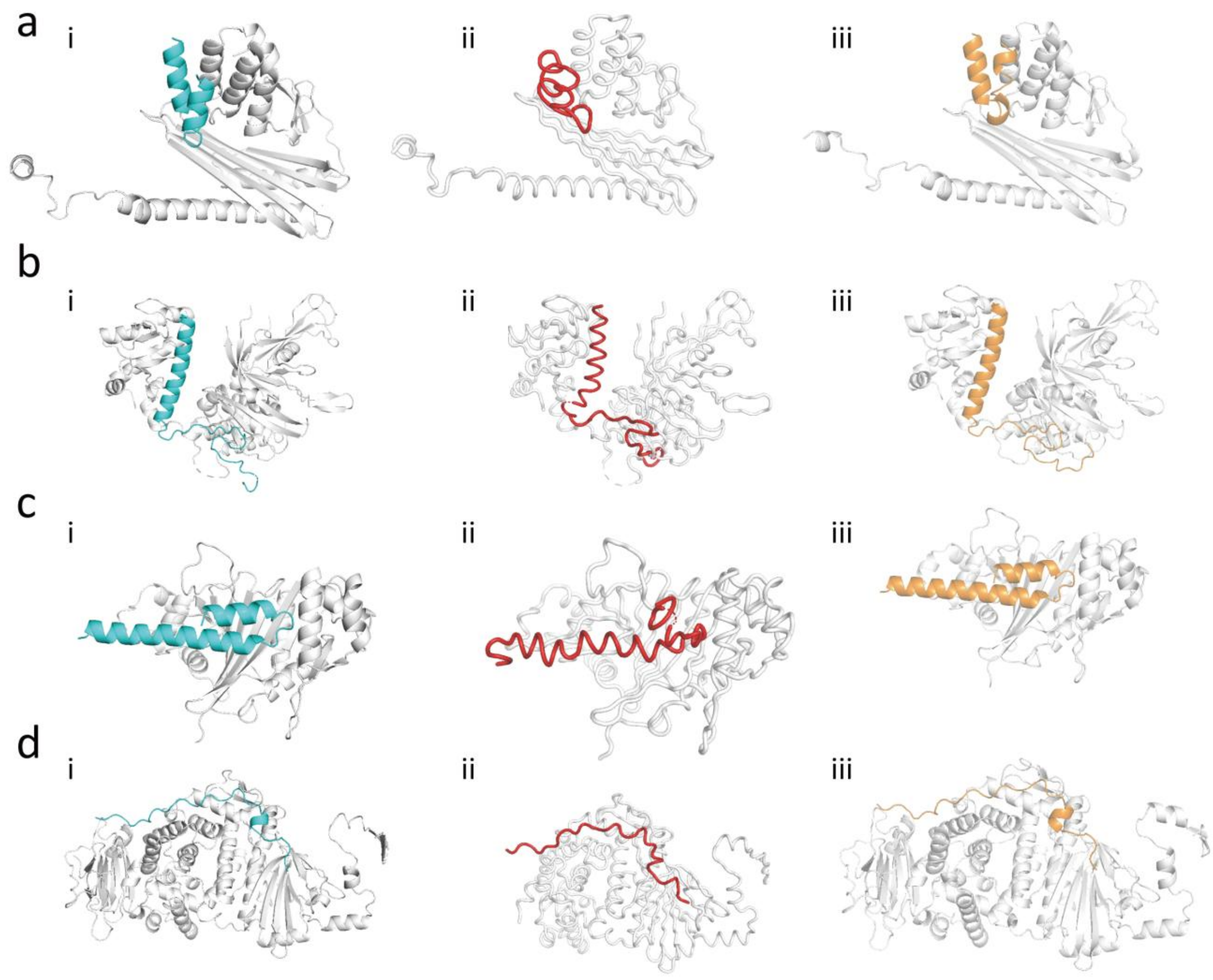


**Supplementary Fig. 28 | Representative structure predictions for long receptor-bound peptides.** a–d, Representative receptor complexes containing long peptide chains with PDB IDs 3LK4, 6AQR, 4W4L and 5TLD, respectively. For each complex, i shows the experimentally determined structure, ii shows the Ourotide prediction and iii shows the AlphaFold 3 prediction. Receptors are shown in white, whereas the experimental, Ourotide-predicted and AlphaFold 3-predicted peptides are shown in cyan, red and orange, respectively. The prediction with the lowest peptide Cα-RMSD among five candidates was selected for each method.

#### Supplementary Tables

#### Supplementary Table 1 | Representative complexes shown in Figs. 2–4.

| PDB ID | Modality | Receptor chain | Ligand chain | Ligand length (aa) | Reported affinity | Affinity index | Regime |
|---|---|---|---|---|---|---|---|
| 1DPJ | Protein–peptide | A | Z | 33 | Ki = 3 nM | 8.5229 | i |
| 1G5J | Protein–peptide | A | Z | 25 | Kd = 0.6 nM | 9.2218 | ii |
| 2PTC | Protein–peptide | A | Z | 58 | Kd = 60 fM | 13.2218 | iii |
| 1MUI | Protein–small molecule | A | Z | — | Ki = 1.3 pM | 11.8861 | — |
| 1A22 | Protein–protein | A | Z | 191 | Kd = 0.34 nM | 9.4685 | — |

Receptor and ligand chains were standardized as A and Z, respectively, for the analyses in this study. The affinity index was calculated as −log10(X [M]), where X is the reported Kd or Ki. Ligand length and recognition regime are not applicable to the small-molecule and protein-interface reference systems where indicated. Affinity annotations were obtained from the PDBbind 2020 index.

#### Supplementary Table 2 | Summary of reference and control datasets.

| Dataset | Source | Selection | Final n |
|---|---|---|---|
| Primary protein–peptide | PDBbind 2020 PP | Ligand chain 8–65 aa; affinity index >7 | 186 |
| Primary protein–protein | PDBbind 2020 PP | Shorter chain >65 aa; affinity index >7 | 246 |
| Primary small molecule | PDBbind 2020 PL and FDA matching | FDA-matched ligand; affinity index >7 | 387 |
| Control protein–peptide | PDBbind 2020 PP | Ligand chain 8–65 aa; no affinity threshold | 349 |
| Control protein–protein | PDBbind 2020 PP | Shorter chain >65 aa; no affinity threshold | 483 |
| Control small molecule | PDBbind 2020 refined set | Structurally eligible; no affinity threshold | 3,055 |

PP and PL denote the protein–protein and protein–ligand subsets of PDBbind 2020, respectively. The affinity index was calculated as $-\log10(X\ [M])$, where X denotes the reported Kd, Ki or IC50. The affinity-unrestricted datasets were used for Supplementary control analyses.

#### Supplementary Table 3 | Physics-guided augmentation acceptance criteria.

| Stage | Descriptor | Acceptance criterion |
| --- | --- | --- |
| Candidate extraction | Peptide length | 8–65 residues |
| Hydrophobic organization | Largest component | ≥3 residues |
| Hydrophobic organization | Second-largest component | ≥3 residues |
| Component energy | Average negative score | $-0.4 < \text{score} < 0$ |
| Component energy | Average positive score | $0 < \text{score} < 3$ |
| Component energy | Positive-score sum | $0 < \text{sum} < 75$ |
| Trunk topology | Trunk length | 3–10 residues |
| Trunk topology | Mean frustration score | <15 |
| Shell organization | S1 − S0 | >−6.5 |
| Shell organization | S2 − S1 | <1.5 |
| Binding-site geometry | Flatness index | >0.8 |
| Final dataset | Retained complexes | 15,325 |

Component-level averages were normalized by the number of residues in the corresponding component. S0 denotes the largest hydrophobic component; S1 and S2 denote successive non-overlapping receptor shells. The flatness index was calculated as $1 - \lambda 3/\lambda 1$ after application of the preceding filters.

#### Supplementary Table 4 | Ourotide training configuration.

| **Setting** | **Structure generator** | **Affinity predictor** |
|---|---|---|
| Representation | Cα geometric graph | Cα geometric graph |
| ESM-2 model | esm2_t12_35M_UR50D | esm2_t12_35M_UR50D |
| Message-passing layers | 4 receptor layers + 6 flow blocks | 4 layers |
| Maximum receptor length | 350 residues | 350 residues |
| Maximum peptide length | 65 residues | 65 residues |
| Optimizer | AdamW | AdamW |
| Main learning rate | $2 \times 10^{-4}$ | $5 \times 10^{-5}$ / $3 \times 10^{-4}$ |
| Candidates per target | 5 | Not applicable |
| Model selection | Validation peptide Cα-RMSD | Validation pKd RMSE |

For the affinity predictor, learning rates of $5 \times 10^{-5}$ and $3 \times 10^{-4}$ were used for the encoder and for the pooling layers and regression heads, respectively. The candidate number refers to independently initialized conformations evaluated for each target at inference.

#### Supplementary Table 5 | Dataset partitions and evaluation sets.

| **Task** | **Dataset or comparison** | **n** |
|---|---|---|
| Structure training | Training set | 12,771 |
| Structure training | Validation set | 1,277 |
| Structure training | Held-out test set | 1,277 |
| Affinity development | Experimental Kd complexes | 213 |
| Affinity development | Receptor clusters | 124 |
| Affinity evaluation | Locked external test set | 49 |
| Structure benchmark | Seven-method Cα-RMSD common set | 47 |
| Structure benchmark | Five-method Cα-RMSD common set | 138 |

The structure-benchmark sample sizes shown here refer to strict common sets used for the Cα-RMSD comparisons. The corresponding valid sample sizes for other metrics are stated with the relevant analysis.

#### Supplementary References


54. Berman, H. M. et al. The Protein Data Bank. *Nucleic Acids Res.* **28**, 235–242 (2000).

55. RDKit: Open-source cheminformatics. https://www.rdkit.org (accessed 10 September 2026).

56. Harris, C. R. et al. Array programming with NumPy. *Nature* **585**, 357–362 (2020).

57. Virtanen, P. et al. SciPy 1.0: fundamental algorithms for scientific computing in Python. *Nature Methods* **17**, 261–272 (2020).

58. Cock, P. J. A. et al. Biopython: freely available Python tools for computational molecular biology and bioinformatics. *Bioinformatics* **25**, 1422–1423 (2009).

59. Hagberg, A. A., Schult, D. A. & Swart, P. J. Exploring network structure, dynamics, and function using NetworkX. In *Proc. 7th Python Sci. Conf.* 11–15 (2008).
